# Coordination of Anle138b to Silver Results in Selective Reduction of a C-terminal truncated Alpha-synuclein Protein and Increased Aggregate Size

**DOI:** 10.1101/2025.10.01.679869

**Authors:** Kelly L. Rue, Susana Herrera, Zhi-Chun Shi, Indranil Chakraborty, Jeremy Tachiki, Josue Ballesteros, Julie K. Andersen, Gordon J. Lithgow, Wisam A. Al Isawi, Gellert Mezei, Minna Schmidt, Raphael G. Raptis

## Abstract

Parkinson’s disease (PD) is a prevalent age-related neurodegenerative syndrome, partially thought to be caused by a decrease in alpha-synuclein proteostasis. Anle138b = 5-(1,3-benzodioxol-5-yl)-3-(3-bromophenyl)-1*H*-pyrazole (**HL**), is undergoing clinical trials as a promising mitigator of alpha-synuclein aggregation. Because complexation to metals is known to modulate the activity of several drugs, we have prepared and characterized: **H_2_L(ClO_4_)**, **[Cu^I^(**µ**-L)]_3_**, and **[Ag^I^(**µ**-L)]_3_**. To better understand the bioviability of these compounds, we monitored their effects in a cell culture model of alpha-synuclein protein aggregation using human alpha-synuclein pre-formed fibrils (PFFs). Using two different anti-alpha-synuclein antibodies, our data suggests that **[Ag^I^(**µ**-L)]_3_** decreases a C-terminal truncated protein that is approximately 12.4 kDa, as well as increases the size and alters the shape of PFF-induced aggregates. This indicates that **[Ag^I^(**µ**-L)]_3_** impacts aggregation in a manner different from **HL** and may serve as a novel tool for studying C-terminal truncation related aggregation chemistry.

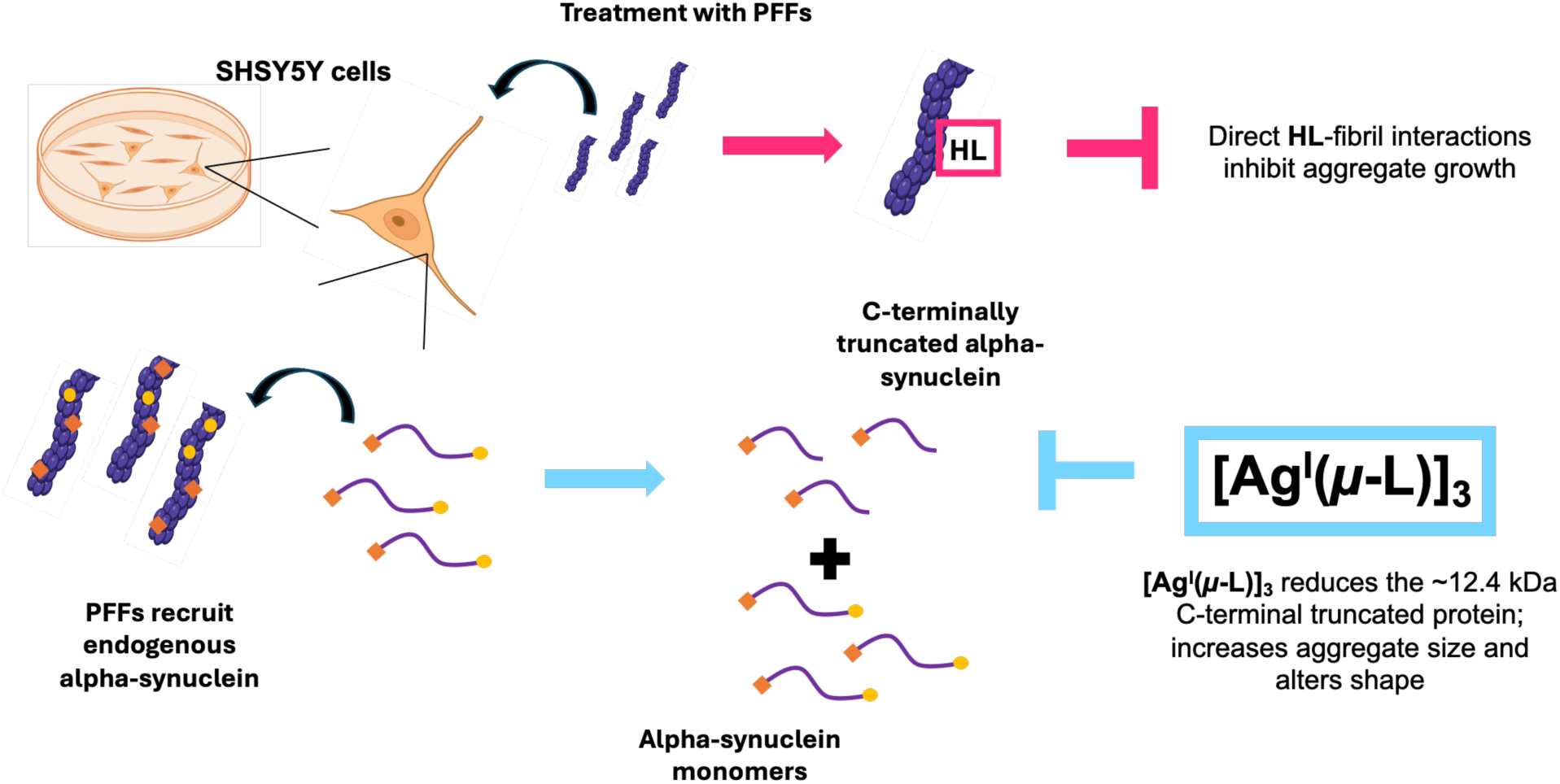

## Introduction

Parkinson’s disease (PD) is a prevalent age-related neurodegenerative syndrome that reduces dopaminergic neuron function and consequentially, results in motor dysfunction (tremor, rigidity, difficulty with balance).^1^ Additionally, people with Parkinson’s (PwPs) often present non-motor symptoms,^2,3^ including gastrointestinal (GI) disturbances, such as constipation, gastrointestinal reflux disease (GERD), and dysbiosis.^4–6^ These symptoms may occur in the prodromal phase and serve as early indicators of PD. Effects on non-motor function have also been associated with the early aggregation of the synaptic protein, alpha-synuclein, and its spread from the GI to the central nervous system.^7–11^ On its own, alpha-synuclein is involved in vesicle transduction and has more recently been found to play an important role in signaling between enteroendocrine and nerve cells.^12–15^ Disruption of alpha-synuclein homeostasis, such as the accumulation of aggregated protein species into toxic oligomers and/or fibrils, can result in the spread of aggregation within and between cells.^16–20^ Furthermore, the etiology of aggregate spread — from the GI versus the olfactory bulb — has recently been correlated with body-first versus brain-first PD.^11,21^ However, the physiological role of alpha-synuclein aggregation is not well understood and while it has been found to be toxic,^22–24^ it has also been suggested to play an alternative role as an anti-microbial peptide (AMPs).^25–27^ Overall, alpha-synuclein aggregates can be either soluble or insoluble^28,29^ and can accumulate into Lewy Bodies, structures found within neurons that may contain other proteins, lipids, and organelle fragments.^30,31^ Importantly, recent breakthroughs have shown that alpha-synuclein aggregation can serve as an important biochemical hallmark of Parkinson’s.^32,33^

More recently, alpha-synuclein C-terminal truncations have been found to play a role in aggregation;^34–39^ several truncated protein species have been identified within Lewy Bodies isolated from PD mouse and human brain samples, as well as tissues from multiple system atrophy (MSA) patients.^40^ Some were found to also interact with tau, a protein implicated in Alzheimer’s disease (AD). Studies have shown that the negatively charged C-terminus plays a protective role against aggregation, repulsing similarly charged alpha-synuclein monomers.^38^ Consequentially, C-terminal truncations have been shown to enhance the rate of protein aggregation and reports have shown that targeting alpha-synuclein at the C-terminus is neuroprotective.^36,39,41^

Currently available medications for Parkinson’s have helped manage symptoms and improve quality of life.^42^ However, most are not known to stop disease progression. A pyrazole-based compound, Anle138b (Anle138b = 5-(1,3-benzodioxol-5-yl)-3-(3-bromophenyl)-1*H*-pyrazole, Scheme 1), which has undergone Phase 1 clinical trials for safety and tolerability in healthy volunteers and those with Parkinson’s,^43,44^ is currently undergoing clinical trials for MSA.^45^ Anle138b is thought to interact with toxic alpha-synuclein oligomers, improving lifespan and lowering protein aggregate load in mouse models of both prion disease and PD. More recently, Anle138b was also shown to interact with alpha-synuclein pre-formed fibrils (PFFs), specifically within the fibril tubular cavity.^46–48^

### Scheme 1

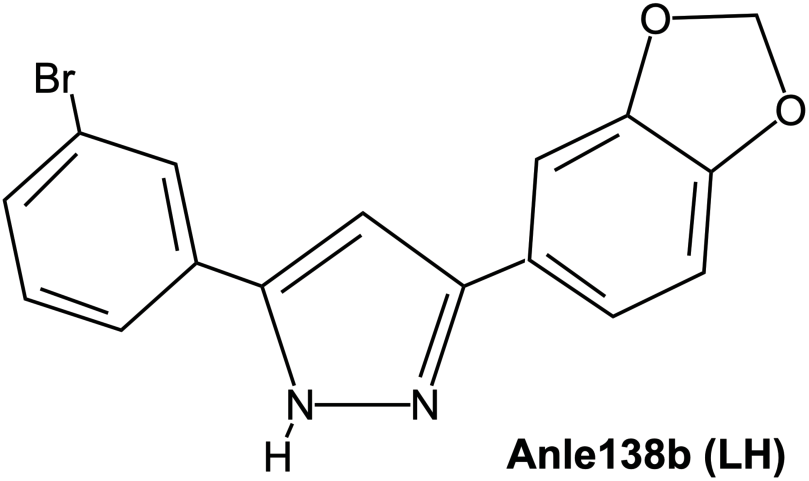

The complexation of between certain drugs and metals has been shown to modulate chemical activity.^49–52^ As the focus of our previous work has been on the complexation and chemistry of transition metals with pyrazole ligands, and pyrazole- and indazole-containing compounds have been investigated for the treatment of PD,^53–55^ we undertook a study of the chemistry and biological viability of several Anle138b metal complexes. First, three new Anle138b-derived materials were synthesized and characterized both structurally and spectroscopically. Then, to better understand their bioviability, treatment was applied to the human derived dopaminergic cell line, SHSY5Y, in an established model of PFF-induced alpha-synuclein protein aggregation.^56^

Here, we present data for three Anle138b **(HL)** derived compounds: **H_2_LClO_4_**, **[Cu^I^(**µ**-L)]_3_**, and **[Ag^I^(**µ**-L)]_3._** Importantly, although copper is known to be a toxic metal substance implicated in Parkinson’s disease etiology,^57^ the stability of these metal-Anle complexes is very strong. We further demonstrate that through western blot analysis, **[Ag^I^(**µ**-L)]_3_** treatment of human differentiated SHSY5Y cells results in a significant reduction of a C-terminally truncated alpha-synuclein protein approximated to be ∼12.4 kDa. This surprising result indicates that the **[Ag^I^(**µ**-L)]_3_** modification causes a unique effect that departs from **HL**. Importantly, we observed that the ∼12.4 kDa protein is C-terminally truncated via experiments which show differential recognition of submonomeric proteins with two different antibodies (BD and CST.) Here, the CST antibody only recognizes residues within the C-terminus and therefore, does not detect the species decreased by **[Ag^I^(**µ**-L)]_3_**. Truncation at the C-terminus is further supported by Edman Sequencing analysis. Differences in antibody detection which correlate with protein modifications, such as C-terminal truncations, have been previously reported.^58^ Our results with confocal imaging further show that treatment with **[Ag^I^(**µ**-L)]_3_** increases the surface area and volume of alpha-synuclein aggregates while decreasing the surface to volume ratio (SVR.) Furthermore, our results show that **[Ag^I^(**µ**-L)]_3_** also alters aggregate shape and increases the preference for the growth over the spread of aggregation. Together, our results project a picture that appears to differ from previous reports which show that inhibition of C-terminally truncated alpha-synuclein proteins reduces aggregation. While we hypothesize that **[Ag^I^(**µ**-L)]_3_** affects protein degradation pathways similar to previous reports,^34^ our results show that treatment increased aggregate size and impacts shape. Future work will explore biochemical mechanisms and track protein aggregation with the help of **[Ag^I^(**µ**-L)]_3_** luminescent properties.

### Experimental section

Chemical reagents and solvents were purchased from Aldrich Chemical Co., Alfa Aesar, Fisher Scientific, and Acros Organics and used as received. Anle138b **(HL)** was prepared using a modified protocol from the literature^59^ and was authenticated by ^1^H-, ^13^C-NMR and HRMS spectroscopic methods (ESI). ^1^H-NMR spectra were recorded on a 400 MHz Bruker Avance spectrometer. FT-IR spectra was recorded with a Perkin Elmer Spectrum 100 FT-IR spectrometer. Mass spectrometric analysis was performed with a Waters Synapt G1 HDMS instrument, using electrospray ionization (ESI) in CH_3_CN solution. Samples were infused by a syringe pump at 5 μL/min and nitrogen was supplied as the nebulizing gas at 500 L/h. The electrospray capillary voltage was set to –2.5 kV, with a desolvation temperature of 110 °C. The sampling and extraction cones were maintained at 40 V and 4.0 V, respectively, at 80 °C. Elemental analyses (CHN) were performed by Galbraith Laboratories Inc. at Knoxville, Tennessee. NMR, FTIR and HRMS spectra can be found in the ESI file.

X-ray diffraction data were collected with a Bruker D8 Quest diffractometer equipped with a PHOTON II detector, using graphite-monochromated Mo-Kα radiation at ambient temperature from single crystals mounted atop glass fibers at random orientation. The APEX3 suite was used for data collection.^60^ Data were corrected for Lorentz and polarization effects.^61^ The structures were solved employing intrinsic phasing with SHELXT and refined by full-matrix least-squares on *F*^2^ with SHELXL.^62,63^ Crystallographic details for **H_2_LClO_4_**, **[Cu^I^(**µ**-L)]_3_**, and **[Ag^I^(**µ**-L)]_3_** are summarized in Table 1.

**Table 1.** X-ray diffraction data and structure refinement parameters.

| | $\text{H}_2\text{LCIO}_4 \cdot (\text{CH}_3)_2\text{CO}$<br>CCDC 2361408 | $[\text{Cu}^{\text{I}}(\mu\text{-L})]_3$<br>CCDC 2361409 | $[\text{Ag}^{\text{I}}(\mu\text{-L})]_3$<br>CCDC 2361410 |
| --- | --- | --- | --- |
| CCDC number | 2361408 | 2361409 | 2361410 |
| Empirical formula | $\text{C}_{19}\text{H}_{18}\text{BrClN}_2\text{O}_7$ | $\text{C}_{48}\text{H}_{30}\text{Br}_3\text{Cu}_3\text{N}_6\text{O}_6$ | $\text{C}_{48}\text{H}_{30}\text{Br}_3\text{Ag}_3\text{N}_6\text{O}_6$ |
| Formula weight | 501.71 | 1217.13 | 1350.12 |
| Temperature/K | 298 | 298 | 172 |
| Crystal system | Triclinic | Triclinic | Triclinic |
| Space group | $P\bar{1}$ | $P\bar{1}$ | $P\bar{1}$ |
| a/Å | 9.6914(4) | 12.1348(4) | 18.0013(8) |
| b/Å | 10.6771(4) | 14.4421(5) | 18.0587(8) |
| c/Å | 10.8957(4) | 14.8078(5) | 18.6632(8) |
| $\alpha/^\circ$ | 99.421(1) | 92.375(1) | 88.76(1) |
| $\beta/^\circ$ | 98.525(1) | 112.941(1) | 99.36(1) |
| $\gamma/^\circ$ | 106.498(1) | 108.701(1) | 62.402(1) |
| Volume/Å<sup>3</sup> | 1043.60(7) | 2222.1(1) | 5224.2(4) |
| Z | 2 | 2 | 4 |
| $\rho_{\text{calc}}/\text{g}\cdot\text{cm}^{-3}$ | 1.597 | 1.819 | 1.717 |
| Reflections<br>collect./indep. | 25526/6286 | 48860/8121 | 89613/17707 |
| Parameters/<sup>a</sup>Obs.<br>Refl. | 339/3705 | 738/8121 | 1331/11714 |
| $R_{\text{int}}$ | 0.0312 | 0.0801 | 0.0528 |
| <sup>b</sup>GooF | 1.033 | 1.045 | 1.027 |
| <sup>c</sup>wR<sub>2</sub> | 0.1443 | 0.2064 | 0.1748 |
| <sup>d</sup>R<sub>1</sub> | 0.0461 | 0.0609 | 0.0608 |

<sup>a</sup>Obs. Refl. = $I_o > 2\sigma(I_o)$

### Synthesis

#### H_2_L(ClO_4_)·H_2_O

(C_16_H_12_BrN_2_O_2_)(ClO_4_)·H_2_O. A mixture of tetrahydrofuran (3 mL) and diethyl ether (6 mL) was added to a vial containing Fe(ClO_4_)_2_·6H_2_O (80.9 mg, 0.32 mmol) and HL (51.6 mg, 0.15 mmol). The mixture was stirred under atmospheric conditions for 30 min, during which a white precipitate formed. The pale, yellow filtrate was removed under reduced pressure. Colorless X-ray grade crystals were obtained via diethyl ether vapor diffusion into acetone. Crystals were washed three times with 5 mL portions of diethyl ether. Anal. Calc. for C_16_H_14_BrN_2_O_7_Cl {(C_16_H_12_BrN_2_O_2_)(ClO_4_)·H_2_O} (461.65 g mol^−1^): C, 41.63; H, 3.06: N, 6.07; Found: C, 41.32; H, 2.77; N, 6.02. IR (*ṽ_max_*, cm^−1^): 3581(w), 3494(w), 3141(w), 2991(w), 1681(m), 1621(m), 1594(m), 1571(m), 1502(m), 1479(s), 1461(s), 1442(s), 1401(w), 1376(w), 1346(w), 1270(w), 1243(s), 1166(w), 1085(s,b), 1029(s), 927(m), 867(m), 838(w), 800(m), 786(s), 738(m), 686(m), 620(m). ^1^H NMR (400 MHz, (CD_3_)_2_SO, 296 K): δ(ppm) = 8.13 (brs, 1H), 8.01 (t, ^3^J = 1.72 Hz, 1H), 7.82 (dt, ^3^J = 2.54, ^2^J = 7.92, 1H), 7.52 (ddd, ^2^J =0.98, ^2^J = 2.92, ^2^J = 7.98, 1H), 7.40 (t, ^3^J = 7.88, 1H), 7.38 (d, ^3^J = 1.60, 1H), 7.33 (dd, ^2^J = 1.72, ^2^J = 8.07, 1H), 7.20 (s, 1H), 7.01 (d, ^2^J = 8.04, 1H), 6.06 (s, 2H).

#### [Cu^I^(µ-L)]_3_

Cu_3_(C_16_H_10_BrN_2_O_2_)_3_. HL (50.0 mg, 0.146 mmol) was dissolved in a 1:1 mixture of THF/MeOH. Then, [Cu(CH_3_CN)_4_]PF_6_ (54.41 mg, 0.146 mmol) and Et_3_N (20 μL, 0.1 mmol) were added with stirring to the solution. The reaction mixture was stirred for 24 h, the white precipitate formed was removed by suction filtration, washed thoroughly with Et_2_O and allowed to dry in a desiccator. The white product was recrystallized by MeOH vapor diffusion into a CH_2_Cl_2_ solution of the product. Elemental analysis for C_48_H_30_Br_3_Cu_3_N_6_O_6_ (1217.14 g mol^−1^), Calcd: C, 47.37; H, 2.48; N, 6.90; Found: C, 47.60; H, 2.41; N, 6.21. FT-IR (*ṽ* _max_, cm^−1^): 2877(w), 2102(w), 1840(w), 1597(m), 1562(w), 1506(m), 1466(vs), 1311(m), 1242(vs), 1211(s), 1116(m), 1036(vs), 930(m), 852(s), 808(m), 769(vs), 746(s), 677(s), 467(m), 409(s). ^1^H NMR (400 MHz, CD_2_Cl_2_, 296 K) δ(ppm) 7.79 (s, 1H), 7.60 (d, ^2^J = 7.60 Hz, 1H), 7.36 (d, ^2^J = 7.76 Hz, 1H), 7.13 (d, ^2^J = 7.96 Hz, 1H), 7.09 (s, 1H), 6.88 (t, ^3^J = 7.74 Hz, 1H), 6.66 (s, 1H), 6.47 (d, ^2^J = 7.88 Hz, 1H), 5.96 (s, 2H).

#### [Ag^I^(µ-L)]_3_

Ag_3_(C_16_H_10_BrN_2_O_2_)_3_. HL (34.3 mg, 0.100 mmol) was dissolved in 6 mL of THF with Et_3_N (14.2 *μ*L, 0.102 mmol) added. AgNO_3_ (16.99 mg, 0.100 mmol) dissolved in 6 mL of MeOH was added to the previous solution and stirred for 6 h protected from light. The solution was filtered and Et_2_O vapors were allowed to slowly diffuse into the colorless filtrate. Colorless crystals were collected and washed with Et_2_O and hexane. Elemental analysis for C_48_H_30_Ag_3_Br_3_N_6_O_6_ (1350.12 g mol^−1^), Calcd.: C, 42.80; H, 2.02; N, 6.24; Found: C, 42.96; H, 1.96; N, 6.13. FT-IR (*v*4_max_, cm^−1^): 3050 (br, w), 3022 (w), 2999 (w), 2886 2999 (br, w), 2760 (w), 1596 (m), 1562 (m), 1506 (s), 1465 (vs), 1395 (w), 1324 (m), 1304 (m), 1239 (vs), 1208 (s), 1150 (w), 1120 (w), 1109 (m), 1073 (w), 1035 (vs), 977 (w), 962 (w), 933 (s), 877 (s), 873 (s), 810 (s), 770 (vs), 742 (s), 702 (w), 692 (w), 679 (s), 620 (w), 598 (w), 497 (w), 478 (m), 466 (w), 422 (w). ^1^H NMR (400 MHz, CD_2_Cl_2_, 293 K): δ (ppm)) 7.79 (s, 1H), 7.57 (d, ^2^J = 7.52 Hz, 1H), 7.43 (d, ^2^J = 7.64 Hz, 1H), 7.26 (d, ^2^J = 8.04 Hz, 1H), 7.11 (s, 1H), 6.98 (t, ^3^J = 8.24 Hz, 1H), 6.68 (s, 1H), 6.57 (d, ^2^J = 7.24 Hz, 1H), 5.97 (s, 2H).

#### PFFs treatment

Type 1 alpha-synuclein pre-formed fibrils (PFFs) were obtained from StressMarq Biosciences. Aliquots were diluted to 1 mg/mL in PBS (VWR) to a final volume of 50 uL and sonicated as per StressMarq protocol to achieve active (shorter) PFFs (Bioruptor 300, 15 minutes HIGH, 30 seconds ON, 30 seconds OFF.) The solution was vortexed and briefly centrifuged with a hand centrifuge prior to addition to cell culture media.

#### Compound solution preparation

All compounds (**HL**, **H_2_L(ClO_4_)**, **[Cu^I^(**µ**-L)]_3_**, and **[Ag^I^(**µ**-L)]_3_**) were initially dissolved in DMSO to a 100 mM stock concentration. Compounds **HL** and **H_2_L(ClO_4_)** dissolved readily in DMSO at a 100 mM stock concentration, forming clear, colorless solutions. **[Cu^I^(**µ**-L)]_3_** and **[Ag^I^(**µ**-L)]_3_** were more difficult to dissolve in DMSO at 100 mM — **[Cu(**µ**-L)]_3_** solution was light green with particulates, **[Ag^I^(**µ**-L)]_3_** solution was a light brown solution with particulates. Initial care was taken to avoid exposure of **[Ag^I^(**µ**-L)]_3_** to light by covering test tubes with aluminum foil. Heat was applied to **[Cu^I^(**µ**-L)]_3_** and **[Ag^I^(**µ**-L)]_3_** solutions and stock concentrations were reduced to 6.25 mM for **[Cu^I^(**µ**-L)]_3_** and 50 mM for **[Ag^I^(**µ**-L)]_3_**. To improve **[Cu^I^(**µ**-L)]_3_** solubility, sonication was initially applied, followed by heating. However, no significant difference was observed in solubility and the solution still appeared somewhat cloudy. Overall, although more soluble than the initial dilution in DMSO at 100 mM, **[Cu^I^(**µ**-L)]_3_** still appeared somewhat cloudy. The turbidity of the **[Ag^I^(**µ**-L)]_3_** solution was more difficult to assess because of its dark color, but upon shaking the solution in the test tube, it appeared homogenous. The solubility for **[Cu^I^(**µ**-L)]_3_** and **[Ag^I^(**µ**-L)]_3_** was improved in cell culture medium for all concentrations except for **[Ag^I^(**µ**-L)]_3_** at 50 and 100 µ M, where precipitation was observed.

#### MTT assay

Non-differentiated SHSY5Y (ATCC) cells were plated into clear 96-well plates at ∼18,000 cells/well. After a 48-hour cell seeding period, cells were treated with vehicle (DMSO), PFFs + vehicle, compound alone, or compound + PFFs for an additional period of 48 hours. Compound toxicity was tested at 6.25, 12.5, 25, 50 and 100 µM concentrations; PFFs treatment was applied at 15 µg/mL (∼1 µM). Cells were initially visualized at the 4, 24 and 48 hr time points to assess any acute effects of toxicity. Dramatic changes in cell morphology and what appeared to be toxic effects were noticed for all compounds at 50 and 100 μM concentration at 4 hours post-treatment. At the 48-hour mark, cells were imaged and then the MTT reagent (MP Bio) was added at 1:10 dilution. Cells were placed back into the cell culture incubator. Incubation times differed for some biological replicates with some being shorter (1.5 hours) and others longer (4 hours.) Cells were observed once again together with MTT reagent. Afterwards, cell culture media was aspirated and 200 uL of DMSO was added directly to the wells. Plates were incubated for 30 minutes in a 37 °C outside the cell culture room. The purple-colored solution in the wells was mixed thoroughly with a multi-channel pipette prior to colorimetric analysis (SpectraMax ABS Plus from Molecular Devices).

#### Western Blot and band quantification

SHSY5Y cells were differentiated for 5 days (3 days of Retinoic Acid (RA) at 10 µM, 2 days of TPA (80 nM)) in 6-well plates at ∼250,000 cells per well, followed by a 48-hour concomitant drug (6.25 µM) and PFFs (5 µg/mL) treatment as described above. After 48 hours, cell culture media was collected and cells were briefly washed with PBS prior to addition of Triton-X (J.T. Baker) soluble buffer (1% Triton-X, 10 mM EDTA in 1X TBS). The cell solutions were scraped using a cell scraper, and pipetted into 1.5 mL Eppendorf tubes, briefly sonicated in the Bioruptor (2 minutes on HIGH, 30 seconds ON, 30 seconds OFF), and centrifuged at 20,000 × g for 20 min. The supernatant (soluble protein) was transferred to a fresh 1.5 mL Eppendorf tube. The pellet was washed with ice cold 1X TBS and centrifuged again at 20,000 x g for 20 minutes and afterwards dissolved in Triton-X insoluble buffer (Triton-X + 2% SDS.) Samples were sonicated until pellet was broken up (minimum of 5 min) with time added accordingly to solubilize the pellet. Protein concentrations were measured via BCA assay (Thermo Scientific) and loaded onto 4-12% gels (NuPage, Invitrogen) for gel electrophoresis (∼80 V for 36 minutes, ∼100 V for 1 hour, ∼120 V for 20 minutes.) Gels were transferred onto a PVDF membrane (Immobilon-pSQ, Millipore) (250 mA, overnight at 4 °C). Both membranes containing soluble and insoluble protein fractions were stained with Ponceau S (Sigma), washed with DI water, and then blocked in a 3% BSA (Sigma), 1X TBST solution for 30 minutes. Blots were incubated with two anti-alpha-synuclein antibodies (BD Transduction Sciences (referred to as BD) and Cell Signaling Technology (CST) as well as anti-actin (Abcam) antibody at 1:1,000 dilution overnight at 4°C. Blots were washed 3 × 10 minutes in 1X TBST, followed by a 1-hour incubation with fluorescent secondary antibodies (Cell Signaling, anti-Mouse IgG DyLight 800 5257P, anti-Rabbit IgG DyLight 680 5366P) at a 1:10,000 dilution. Blots were washed again for 3 × 10 minutes in 1X TBST (TBS from VWR, Tween 20 VWR) and then visualized using a fluorescent scanner (Odyssey LI-COR.) All band intensities were quantified using Fiji or ImageJ. Relative molecular weights of bands mentioned were quantified in ImageJ with protein ladder (BioRad) as reference.

#### Immunocytochemistry (ICC)

Non-differentiated SHSY5Y cells were plated at cell density 54,000 cells/well into confocal slides (Ibidi, 15 µ-Slide 8 well High ibiTreat). Cells were grown for 7 days total and PFFs + compound were added from day 5 to day 7 at a concentration of 5 µg/mL. To avoid complete extraction of cell culture medium and exposing cells to air, addition of solution was performed sequentially removing 100 µL of cell culture media, followed by addition of 100 µL of treatment solution. This was repeated 4-5 times in the four different corners of the well, plus the middle of the well, to ensure cell culture media was replaced with compound solution. On the last day of the experiment, cells were fixed in 4% PFA solution for ∼15 min. PFA (Chem Cruz) was added sequentially to avoid exposure to air. First, 250 µL of PFA was added, followed by removal of 250 µL of PFA + cell culture media × 4, to ensure complete replacement of cell culture media with PFA. Cells were washed with ice-cold PBS 3 × 5 min and stored in PBS at 4°C until ICC. After fixation, cells were permeabilized with 0.5% Triton-X in 10% Natural Goat Serum (NGS) for 40 min, followed by blocking for 20 min with 10% NGS. Primary ICC cocktails were prepared in 1% NGS (Jackson Immuno Research). BD antibody was diluted 1:200. Cells were incubated in primary antibody for ∼48 hrs. at 4°C followed by secondary incubation at 1:1,000 dilution for 1 hr. at room temperature. Nuclei were stained with DAPI at 1:1,000 for 10 min at room temperature. Z-stack images were acquired using a Zeiss LSM 980 microscope with Airycan2 and surface area, volume, puncta number, prolate, oblate, and sphericity quantifications were measured using 3D objects in Imaris (RRID:SCR_007370).

#### Edman Sequencing

Non-differentiated SHSY5Y cells were treated with PFFs. Western blot samples were prepared as described above. Gel electrophoresis was performed on a 10% gel in MOPS buffer. Because the blot required marking of the Ponceau S stained bands, only the insoluble fraction was analyzed as it solely contained enough protein to be visualized with Ponceau S. Four technical replicates were performed and the bands were pooled together from the replicates i.e. according to their horizontal position. The blot was shipped and then loaded onto the sequencer (Instrument: Shimazu PPSQ-53A, Column: Wakopak Wakosil PTH-GR, Solvents: Wako PTH-amino acids Mobile Phase A, Wako PTH-amino acids Mobile Phase B, Flow: 300 µL/minute, Gradient elution, Detection: 269 nm).

#### Statistical analysis

Statistical analysis was performed on all biological replicates using Prism 10 software or ChatGPT generated R programming language. In analysis of MTT results, sum normalization and y-to-y^2^ transformation (y = y^2, in Prism 10 shorthand notation) was applied prior to analysis with 2-Way ANOVA Tukey multiple comparisons test. For western blot samples, results were pooled between blots with PFFs, **HL**, **H_2_LClO_4_**, and **[Cu^I^(**µ**-L)]_3_** treated cells, while, due to space constrains, separate blots were run with PFFs and **[Ag^I^(**µ**-L)]_3_** only. Data was transformed using the y-to-y^1/2^ function (y = √y, in Prism 10 shorthand notation) followed by 2-Way ANOVA with Tukey’s multiple comparisons test. For confocal microscopy, up to 10 images were taken for each condition. Individual puncta were analyzed with generation of masks/objects in Imaris software. The following data was collected: surface area, volume, ellipticity (prolate and oblate values), and sphericity, as well as puncta and nuclei number. While the original Anle138b compound (**HL**) and **H_2_LClO_4_** were also visualized, puncta distinction was difficult to achieve, resulting in skewed results. Therefore, analysis included only visualization of **[Cu^I^(**µ**-L)]_3_** and **[Ag^I^(**µ**-L)]_3_** related puncta. Data handling and statistical analysis was performed in R studio with code generated through ChatGPT interface. To determine statistical significance, data was fit to a linear mixed-effects model to account for differences between biological replicates and between puncta within replicates. Here, each treatment had a fixed effect, while a random intercept was found for each replicate to account for between-replicate baseline shifts and for within-replicate clustering. Then, the estimated marginal means (EMMs) were determined where values were given equal weights (emmeans). The log-transformed outcomes in EMMs were back-transformed to the original scale. Dunnett’s adjustment was applied to compare each treatment to control (PFFs). Count-size tradeoff: Incidence Rate Ratio (IRR) was calculated by dividing the mean of the puncta/nucleus in the treatment group by the mean of the puncta/nuclei in the control group; Geometric Mean Ratio (GMR) was calculated by dividing the geometric mean of the treatment group by the geometric mean of the control group; Mass balance: Total Volume per Nuclei Ratio (TVNR) = (sum of the puncta volume in the treatment group)/(the number of nuclei for the treatment group) divided by (sum of the puncta volume in the control group)/(the number of nuclei for the control group). Omnibus ANCOVA with Satterthwaite degrees of freedom was applied to determine differences among the treatment groups; mixed-effects ANCOVA with Satterthwaite t-tests were applied to determine β (common slope, differences from zero); pair-wise Tukey adjustment was applied to determine Δβ (differences between treatment groups) (Table 1.)

## Results

**H_2_LClO_4_** crystallizes in the triclinic *P*1̅ space group with the entire molecule in the asymmetric unit (Figure 1). The positive charge of the H_2_L ion is balanced by a perchlorate anion. This also helps to stabilize the crystal structure, along with an acetone molecule, through N(H)···O with donor-acceptor separations hydrogen bonds of 2.743(4) Å and 2.917(5) Å, respectively. Two oxygen atoms of the perchlorate ion are disordered (Figure S5) over two positions; the other two are ordered due to intermolecular interactions. The pyrazolium ion is almost perfectly planar with a dihedral angle of 6.27° between the bromophenyl and the benzodioxole planes. The benzodioxole group is slightly more twisted than the bromophenyl group in relation to the pyrazole plane, with dihedral angles of 7.80° and 3.43°, respectively. A crystal packing diagram shows that the H_2_L ions form channels, with an inter-channel distance ranging between 4.5 Å and 5.5 Å, where the perchlorate and acetone molecules reside (Figure S5).

**Figure 1.**
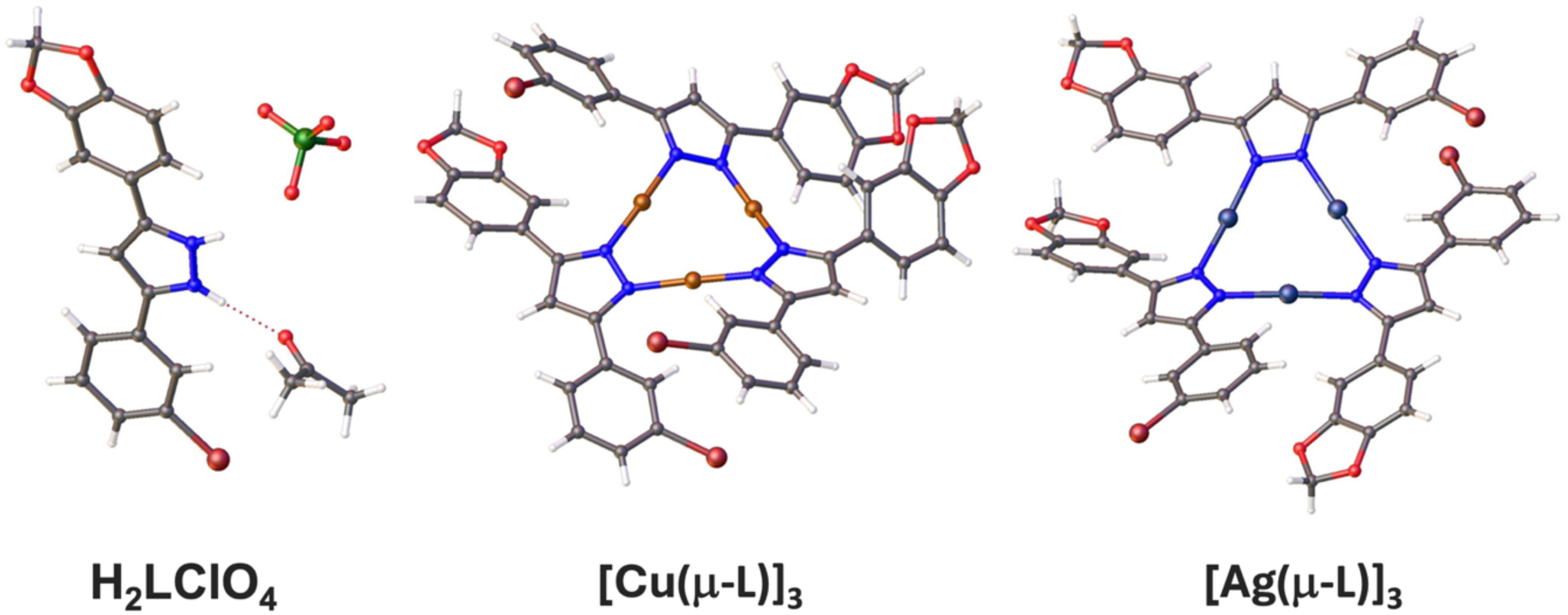
Ball-and-stick diagram of **H_2_L(ClO_4_)**·(CH_3_)_2_CO, **[Cu^I^(**µ**-L)]_3_** and **[Ag^I^(**µ**-L)]_3_** (crystallographic disorder of the perchlorate anion and phenyl groups not shown).

Both **[Cu^I^(**µ**-L)]_3_** and **[Ag^I^(**µ**-L)]_3_** crystallized in the triclinic space group *P*1̅ with one and two whole molecules in the asymmetric unit, respectively, and the 3- and 5-substituents of the pyrazolate ring crystallographically disordered, as described in the Supporting Information. The Cu-centers are in an approximately linear two-coordinate environment, with Cu – N bonds between 1.77(3) Å and 1.92(2) Å, whereas and Ag – N bond lengths are approximately 2.080(2) Å. The nine membered Cu_3_N_6_ metallacycle is significantly distorted from ideally planar geometry, with intra-trimer Cu^…^Cu distances of 3.245 Å, 3.161 Å, and 3.071 Å and a torsion angle of the metallacycle plane of 0.258° (Figure 1); the corresponding Ag^…^Ag distances range from 3.312 Å to 3.405 Å. The shortest inter-trimer contacts between adjacent Cu_3_ or Ag_3_ units are 4.415 Å and 4.175 Å, respectively (Figure S6, S7).

### UV-Vis/Fluorescence

The UV-vis absorption spectra of **HL**, **H_2_L(ClO_4_)**, **[Cu^I^(**µ**-L)]_3_** and **[Ag^I^(**µ**-L)]_3_** (Figure S8) consist of features in the area < 350 nm, whereas the emission spectra of **HL**, **H_2_L(ClO_4_)** and **[Cu^I^(**µ**-L)]_3_** show maxima at 350 and 680 nm, in addition to a peak at 480 nm of **H_2_L(ClO_4_)** (Figure S9). However, the emission spectrum of **[Ag^I^(**µ**-L)]_3_** (Figure 2) shows an intense emission at 460 nm, differentiating it from **HL** and the other two compounds and consistent with literature reports of luminescent properties of polynuclear Ag^I^ complexes.^64,65^ In the solid state, **[Ag^I^(**µ**-L)]_3_** luminesces with a λ_max_ = 450 nm (Figure S10).

**Figure 2.**
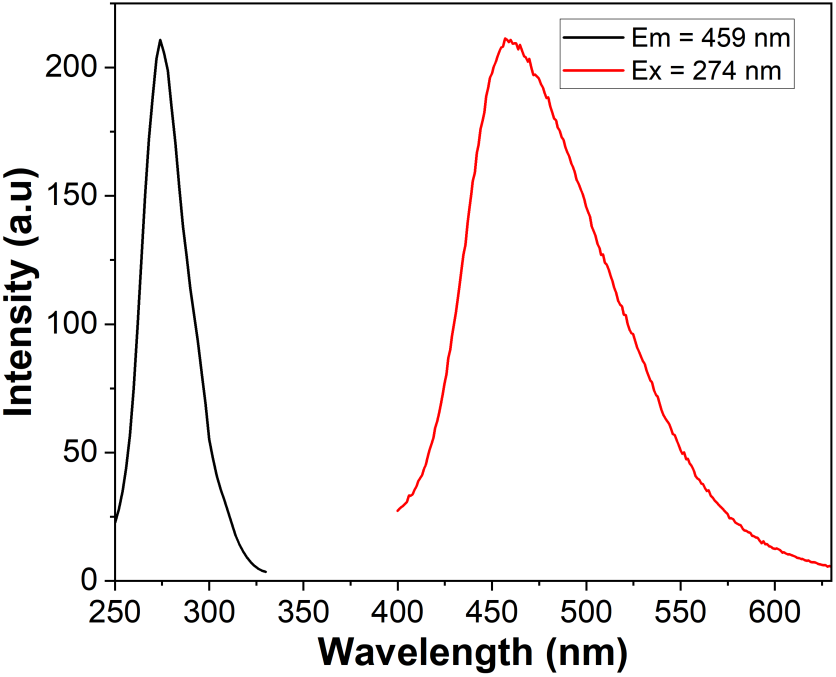
UV-Vis emission and excitation spectra of 0.05 M MeOH/thf solution of **[Ag^I^(**µ**-L)]_3_**, 295 K.

### MTT results show similar bioviability for HL, H_2_LClO_4_, [Cu^I^(µ-L)]_3_ and [Ag^I^(µ-L)]_3_ at 6.25 mM concentrations

The bioviability of **HL**, **H_2_LClO_4_**, **[Cu^I^(**µ**-L)]_3_,** and **[Ag^I^(**µ**-L)]_3_** was assessed using the MTT cell viability assay in non-differentiated human dopaminergic SHSY5Y cells. In this assay, 3-(4,5-Dimethylthiazol-2-yl)-2,5-Diphenyltetrazolium Bromide is converted to a purple formazan crystal within mitochondria. While argued to be an assessment of mitochondrial health more than cellular viability,^66^ we find that in our hands, the MTT assays can be the most consistent test for determining chemical bioviability. In addition, alpha-synuclein PFFs have been observed to target mitochondria.^67^ We therefore utilized the MTT assay to assess whether the compounds exert protective effects against a moderately toxic PFFs dose — 15 µg/mL or ∼1 µM — by treating cells with the following conditions over a 48-hour period: 1) vehicle (DMSO), 2) PFFs + vehicle, 3) compound at the following concentrations: 6.25, 12.5, 25, 50, or 100 µM, and 4) concomitant treatment of PFFs + compound. Our results indicate that the PFFs dose chosen reduced MTT signal by 20-30% (Figure 3). In general, we found that all compounds exhibited no significant toxicity at 6.25 µM and 12.5 µM.

**Figure 3.**
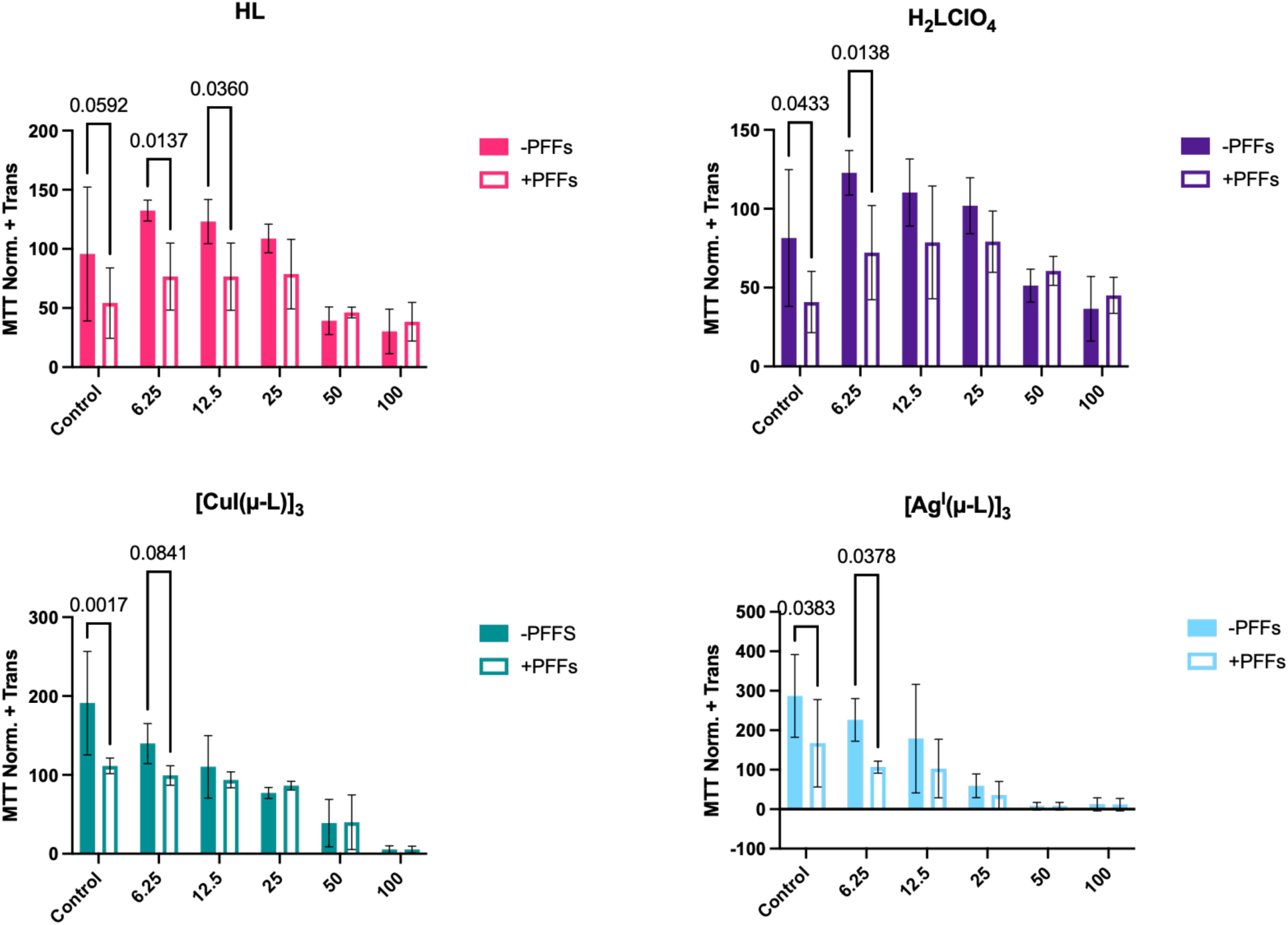
MTT assay results for compounds **HL**, **H_2_LClO_4_**, **[Cu^I^(**µ**-L)]_3_**, and **[Ag^I^(**µ**-L)]_3_** in absence or presence of PFFs. All compounds show no toxicity at 6.25 µM and 12.5 µM doses. A slight patterned increase in MTT values is present between -PFFs and +PFFs for **HL** and **H_2_LClO_4_** at concentrations 50 and 100 µM, but these differences were not significant (p = 0.7415, p = 0.6985 for **HL** and p = 0.6291, p = 0.6559 **H_2_LClO_4_** for 50 and 100 µM, respectively.) **[Cu^I^(**µ**-L)]_3_**-PFFs exhibited toxicity at 25 µM but with +PFFs, only showed toxicity at 50 µM (p ≤ 0.0004 and p ≤ 0.0409, respectively.) There is a slight but non-significant difference between -PFFs and + PFFs at concentration 25 µM (p = 0.6850.) **[Ag^I^(**µ**-L)]_3_** shows toxicity starting at the 25 µM for -PFFs (p ≤ 0.0041.) (Sum normalization with y = y^2 transformation, 2-Way ANOVA and Tukey’s multiple comparisons test; n = 3.)

For almost all compounds, addition of PFFs to chemical doses of 6.25 and 12.5 µM also decreased MTT values in comparison to compound-alone conditions. However, at the 25 µM dose, the differences between cells with or without PFFs appeared to lessen.

### [Ag^I^(µ-L)]_3_ treatment decreases ∼12.4 kDa C-terminally truncated alpha-synuclein protein in both soluble and insoluble protein fractions

To determine the effect of compound treatment upon alpha-synuclein protein aggregation, we treated cells with a lower PFFs dose — 5 µg/mL. This dose was previously determined to yield alpha-synuclein aggregate growth over a 48-hr. treatment period (data not shown.) Using the concomitant method of treatment as described in the MTT assay, differentiated SHSY5Y cells were treated with compound alone or with compound plus PFFs. After 48 hrs., proteins were extracted with Triton-X soluble and Triton-X insoluble and alpha-synuclein was visualized using an anti-alpha-synuclein antibody from BD Transduction Sciences that recognizes residues 15-123 (Figure 4). Here, we see that PFFs treatment shows a significant increase in total alpha-synuclein signal (Figure S11). While alpha-synuclein is a 140 kDa protein, it is known to appear at ∼19 kDa. We see in our blots it appears at ∼17 kDa, which is both blots — -PFFs and + PFFs (Figure 4, red arrow, left and right panels.) We also observed a 15 kDa band (blue arrow), for which signal was noticeably increased. In addition, PFFs addition resulted in strong signal of several bands below the monomer, indicating truncation of alpha-synuclein protein. We also observed that **[Ag^I^(**µ**-L)]_3_** treatment reduced a very specific band in the soluble and insoluble fractions (Figure 4, orange arrow.) After estimating molecular weights with ImageJ software (Figure S12), we found that this band is ∼12.4 kDa with some variation. Specifically, in the blots which only contain PFFS and **[Ag^I^(**µ**-L)]_3,_** we observed that this band is at ∼12.4 kDa in the insoluble fraction while is at ∼12.2 kDa in the soluble fraction (Figure S12 C,D). Due to limitation in space, samples for **HL**, **H_2_LClO_4_**, and **[Cu^I^(**µ**-L)]_3_** were run on a separate gel. Here, we observed the band we approximate to be decreased by **[Ag^I^(**µ**-L)]_3_** at ∼12.6 kDa in the soluble fraction, but at ∼11.6 kDa in the insoluble fraction (Figure S12 A, B.) For ease of reference, we averaged the values from the blots which contain PFFs and **[Ag^I^(**µ**-L)]_3_** only for an approximately molecular weight of 12.4 kDa. Future experiments will explore exact molecular weight with mass spectrometry.

**Figure 4.**
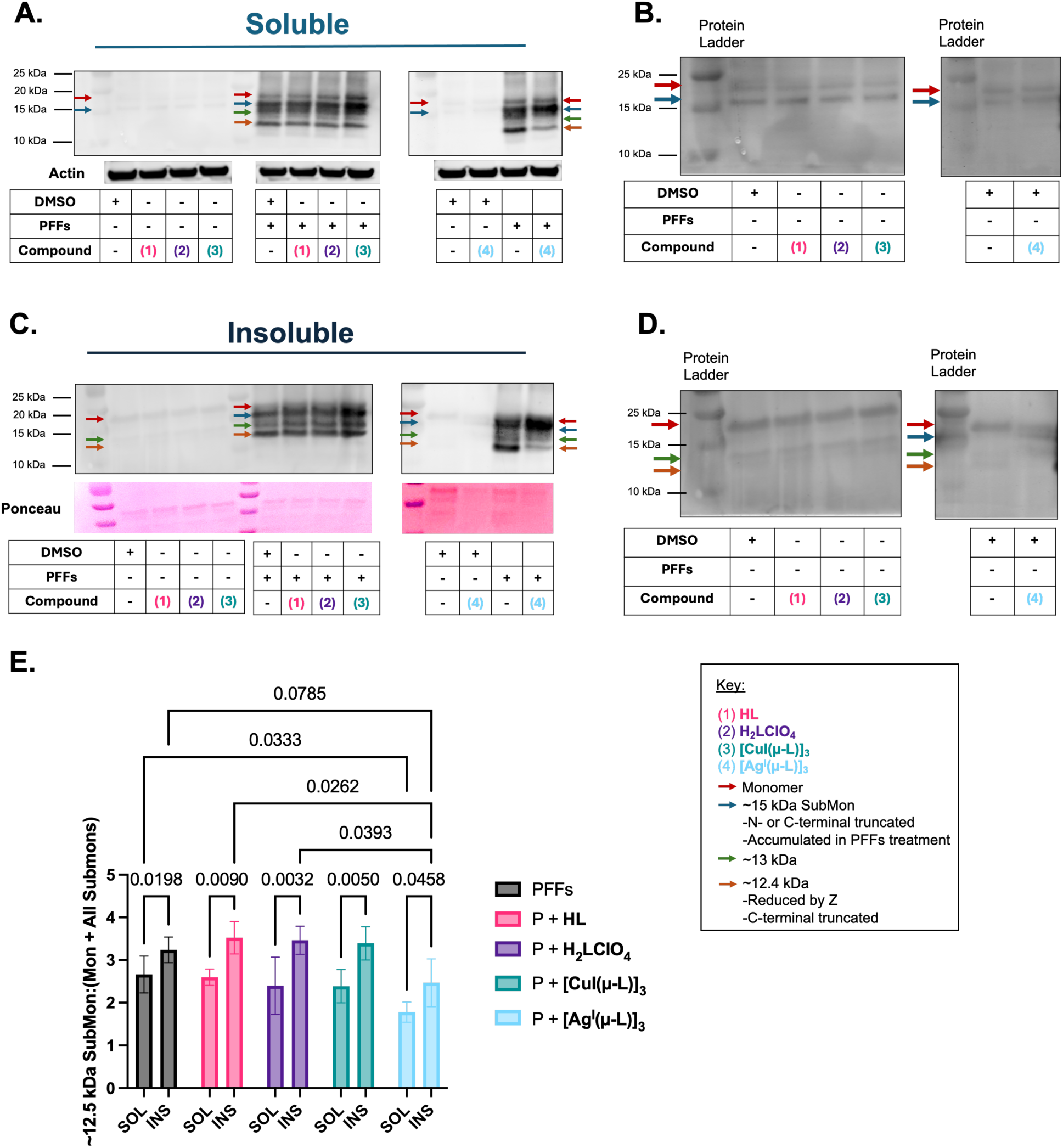
Western blot results showing effect of compounds **HL** (1), **H_2_LClO_4_** (2), **[Cu^I^(**µ**-L)]_3_** (3), and **[Ag^I^(**µ**-L)]_3_** (4) on alpha-synuclein monomer and submonomer species in the soluble and insoluble fractions +/-PFFs (P). **(A, C)** Representative western blot images of soluble and insoluble fractions, respectively, show marked reduction of ∼12.4 kDa submonomer with **[Ag^I^(**µ**-L)]_3_** treatment as compared to control and **HL**, **H_2_LClO_4_**, and **[Cu^I^(**µ**-L)]_3_**. **(B, D)** Enhanced western blot images of control cells (-P) of soluble and insoluble fractions, respectively. Images show presence of the monomer (∼17 kDa) (red arrow) and truncated species ∼15 kDa (blue arrow) in the soluble fraction, and increased quantities of the monomer (∼17 kDa, red arrow), ∼15 kDa (blue arrow), ∼13 kDa (green arrow), and ∼12.4 kDa species (orange arrow) in the insoluble fraction. **(E)** Quantification of the ∼12.4 kDa band with respect to the total signal observed from the monomer plus all submonomers shows significantly more of this band in the insoluble (INS) fraction as compared to the soluble (SOL) fraction for all compounds. There is also a significant, or trending decrease for this species in samples treated with **[Ag^I^(**µ**-L)]_3_** in the SOL and INS fractions (p ≤ 0.0333 and p ≤ 0.0785), respectively (2-Way ANOVA, Tukey multiple comparisons test, n = 3).

In order to quantify the difference in the ∼12.4 kDa species due to **[Ag^I^(**µ**-L)]_3_** treatment, we utilized an internal method of protein quantification and normalization by taking the ratio of the ∼12.4 kDa submonomer to the total signal from the monomer plus all submonomers (Mon + All SubMons, Figure 4). This was done in order to sidestep difficulty with protein normalization in insoluble fractions (Ponceau S staining was used as a reference but not utilized for normalization; actin levels were measured but treatment conditions in the insoluble fraction may alter these protein levels (Figure S11)). In comparing these ratios, we observed that **[Ag^I^(**µ**-L)]_3_** significantly decreased the ∼12.4 kDa band in the soluble fraction (p ≤ 0.0333, Figure 4 A, E) and showed a significantly trending effect in the insoluble fraction (p = 0.0785) (Figure 4 C, E).

Using a best practice method, we treated these blots with an additional alpha-synuclein antibody which recognizes only residues located within the C-terminus (CST, surrounding proline 128 and including residues 122 and 123), and which we previously noted does not detect some of the same submonomeric species (not shown.) Here, we noticed that the submonomeric species, including the ∼12.4 kDa species, were not detected in either the soluble or in the insoluble fractions with the CST antibody (Figure 5 A,C, orange arrow). Given that the BD antibody recognizes a greater segment of the alpha-synuclein protein, including residues in the N-terminus, while the CST antibody recognizes residues only within the C-terminus, these results suggest that the ∼12.4 kDa decreased by **[Ag^I^(**µ**-L)]_3_** treatment is C-terminally, and not N-terminally, truncated. This result was further corroborated by Edman Sequencing analysis of the insoluble fraction (Figure S13.) While both the soluble and insoluble fractions were extracted, only the insoluble fraction showed enough protein upon Ponceau S staining (visualization of bands are required for loading the proteins onto the sequencer (Figure S13B.)) However, as exact determination of the bands is not possible because Edman Sequencing does not involve antibody staining, the bands were labeled ‘top’, ‘middle’, and ‘bottom’ for proteins likely corresponding to the monomer, ∼15kDa and ∼12.4/13kDa bands, respectively. The order and visualization of the proteins matched our previous observations as well (Figures 4, S11, S13.) Altogether, the sequencing results showed that the middle and bottom bands signified the presence of the MDVFM motif, which includes the first five amino acids of the alpha-synuclein protein sequence beginning from the N-terminus. The top band however, only showed the DVF motif where the lack of the other two amino acids suggested that the strength of the signal was subdued due to signal from other proteins in the sample. Altogether, presence of the MDVFM motif in the middle and bottom bands, together with the differences in detection between the BD and CST antibodies, suggest that alpha-synuclein is truncated at the C- and not the N-terminus and that the ∼12.4 kDa is a C-terminally truncated alpha-synuclein protein.

**Figure 5.**
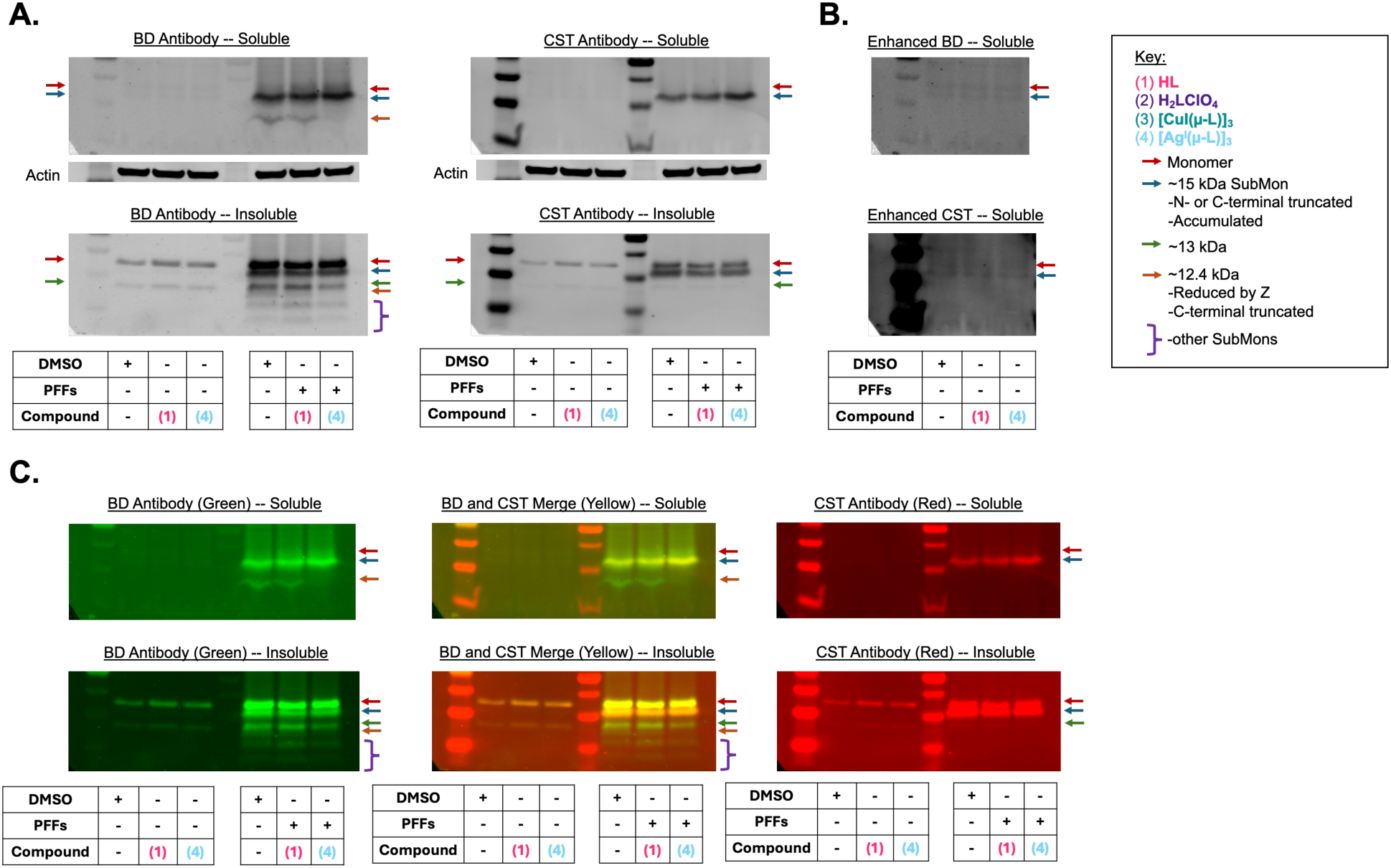
**(A-B)** Representative western blot images from non-differentiated SHSY5Y cells show differences between detection patterns of the BD antibody (recognizing residues 15-123) and the CST antibody (recognizing residues in the C-terminal domain surrounding proline 129, and including residues 122 and 123) as detected by fluorescent secondary antibodies. The BD antibody shows strong detection of the ∼12.4 kDa band (orange arrow) in both fractions, while the CST antibody shows no visible detection in the soluble fraction, and little visibility in the insoluble fraction. The differential detection patterns of the BD and CST antibodies indicate that the ∼12.4 kDa protein is C-terminally truncated. **(C)** Overlay of fluorescent secondary antibodies confirms mutual detection of the ∼15 kDa species (blue arrow) by both BD and CST antibodies in the soluble and insoluble fractions. Detection of the monomer (red arrow) is greater in the insoluble fraction for both antibodies showing strong overlap in the middle image representing the overlay of both antibodies. The BD antibody (green blot) strongly detects the ∼12.4 kDa band in the soluble and insoluble fraction. However, this band is not detected by the CST antibody (red blot). Slight detection is present in the insoluble fraction upon visualization with the CST antibody.

### Confocal microscopy shows [Ag^I^(µ-L)]_3_ results in larger and differently shaped aggregates, but does not promote aggregate spread

Further imaging with confocal microscopy showed that alpha-synuclein aggregates clearly form within non-differentiated SHSY5Y cells upon treatment with PFFs (Figure 6A). Here, the endogenous alpha-synuclein (red), which is normally spread over the nucleus and surrounding cellular area (control, left panels), is gathered into puncta upon addition of PFFs (right panel.) In order to better understand the effect of **HL**, **H_2_LClO_4_**, **[Cu^I^(**µ**-L)]_3_**, and **[Ag^I^(**µ**-L)]_3_** on alpha-synuclein aggregation, we characterized the individual aggregate sizes (surface area, volume, and surface to volume ratio (SVR)) as well as the shape (prolate, oblate, and sphericity) using 3D surfaces generated in the Imaris program. Aggregate numbers ranged from several hundred individual puncta to over a thousand. A similar method was recently used to characterize A-beta plaque formation in mouse brain tissues.^68^ However, upon analyzing aggregates from **HL** and **H_2_LClO_4_** treated cells, we saw that size and shape characterization was difficult to achieve in these treatments due to lower definition of puncta structure — although alpha-synuclein signal does show puncta formation, it remains spread out, indicating that the compounds abrogate the PFFs-induced aggregation as previously published for **HL** (Figure S14). For this reason, we limited our method of characterization to the puncta formed with **[Cu^I^(**µ**-L)]_3_** and **[Ag^I^(**µ**-L)]_3_** treatment.

**Figure 6.**
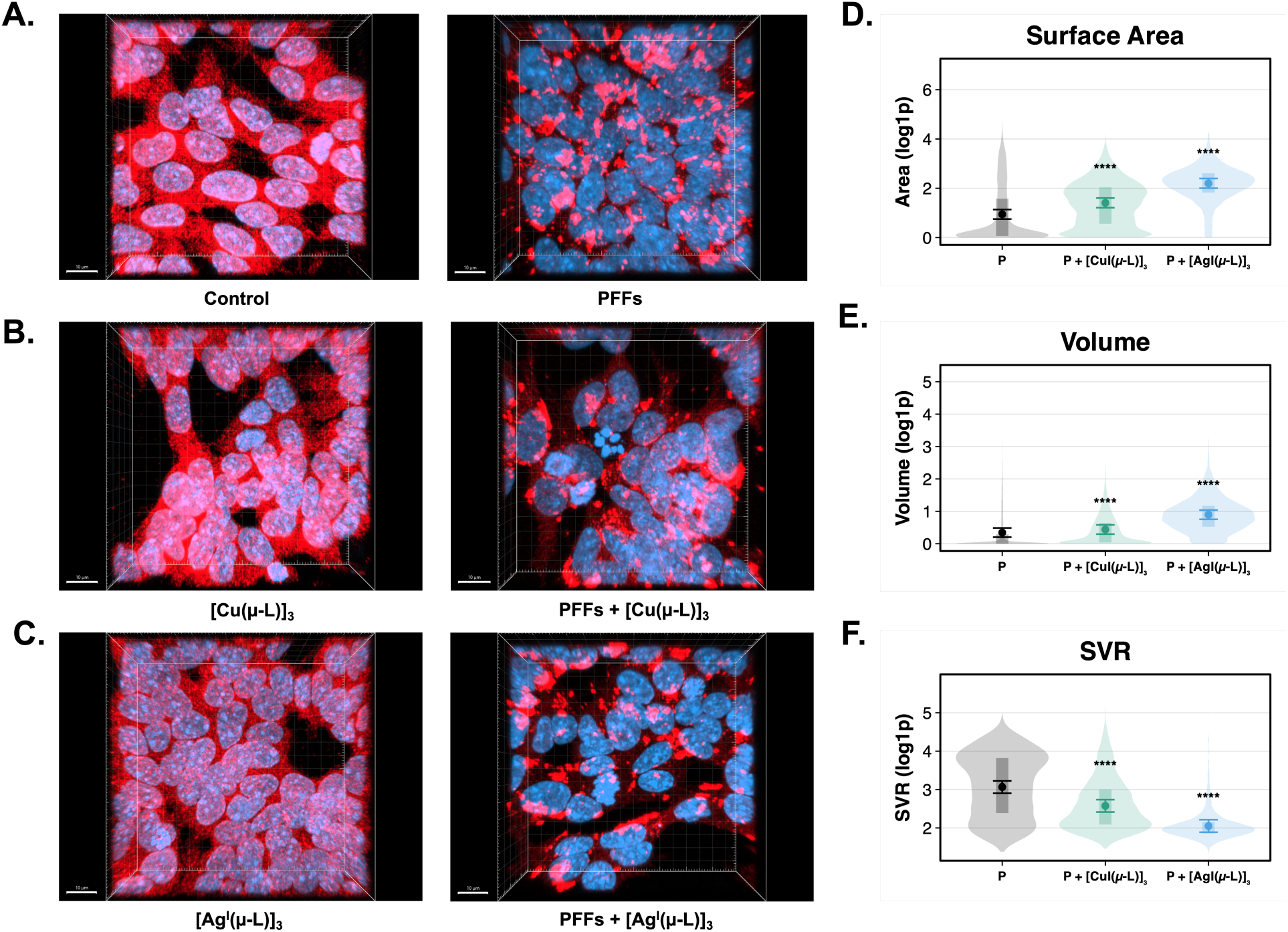
Representative confocal images and characterization of alpha-synuclein aggregates (red) formed due to PFFs treatment in SHSY5Y cells. **(A-C) [Cu^I^(**µ**-L)]_3_** and **[Ag^I^(**µ**-L)]_3_** treatment results in aggregates of larger sizes and altered shapes as compared to PFFs (P) only treatment. **(D-F)** Quantification and comparison of individual puncta show significant differences in surface area, volume, and surface to volume ratio (SVR), with the strongest differences present in **[Ag^I^(**µ**-L)]_3_** treated cells. Mixed-effects model with natural log transform showed group differences in surface area and volume. Back transformed EMMs (mean [95% CI]) were: PFFs area 0.94 [0.75-1.14] µm^2, volume 0.34 [0.20-0.49] µm^3), and SVR 3.06 [2.90-3.22]. **[Cu^I^(**µ**-L)]_3_** increases surface area to 1.41 [1.21-1.60] µm^2, slightly increases volume 0.44 [0.30-0.58] µm^3 and decreased SVR to 2.58 [2.41-2.74]. **[Ag^I^(**µ**-L)]_3_** shows the largest increase in both surface area 2.20 [2.00-2.40] µm^2 and volume 0.90 [0.75-1.04] µm^3 while substantially decreasing SVR to 2.05 [1.89-2.21]. Relative to PFFs, the Dunnett-adjusted ratio with 95% CI for surface area, volume and SVR for **[Cu^I^(**µ**-L)]_3_** is: 1.59 [1.55, 1.64], 1.10 [1.08, 1.12], and 0.61 [0.60-0.63], respectively, and for **[Ag^I^(**µ**-L)]_3_**: is 3.51 [3.36, 3.67], 1.74 [1.70, 1.78], and 0.36 [0.35-0.37], respectively (n=3, ****p < 0.0001)).

Here, we saw that PFFs treatment resulted in aggregates with an average surface area of 0.94 µm^2, a volume of 0.34 µm^3, and an SVR of 3.06. **[Cu^I^(**µ**-L)]_3_** treated cells show a significantly higher average surface area (1.41 µm^2) and volume (0.44 µm^3), while the SVR decreased to 2.58 (****p<0.0001). **[Ag^I^(**µ**-L)]_3_** however, showed the largest difference: average surface area was measured at 2.20 µm^2, volume at 0.90 µm^3, and SVR was reduced to 2.05 (Figure 7D-F) (****p<0.0001). Importantly, these results also show that **[Ag^I^(**µ**-L)]_3_** treatment leads to more uniform puncta as compared to both PFFs and **[Cu^I^(**µ**-L)]_3_**, which can be seen from the smaller degree of variation present within the violin plots. Altogether, our data shows that while both metal chelated compounds significantly increase aggregate size but decrease SVR, the largest and most consistent aggregates are formed with **[Ag^I^(**µ**-L)]_3_** treatment.

**Figure 7.**
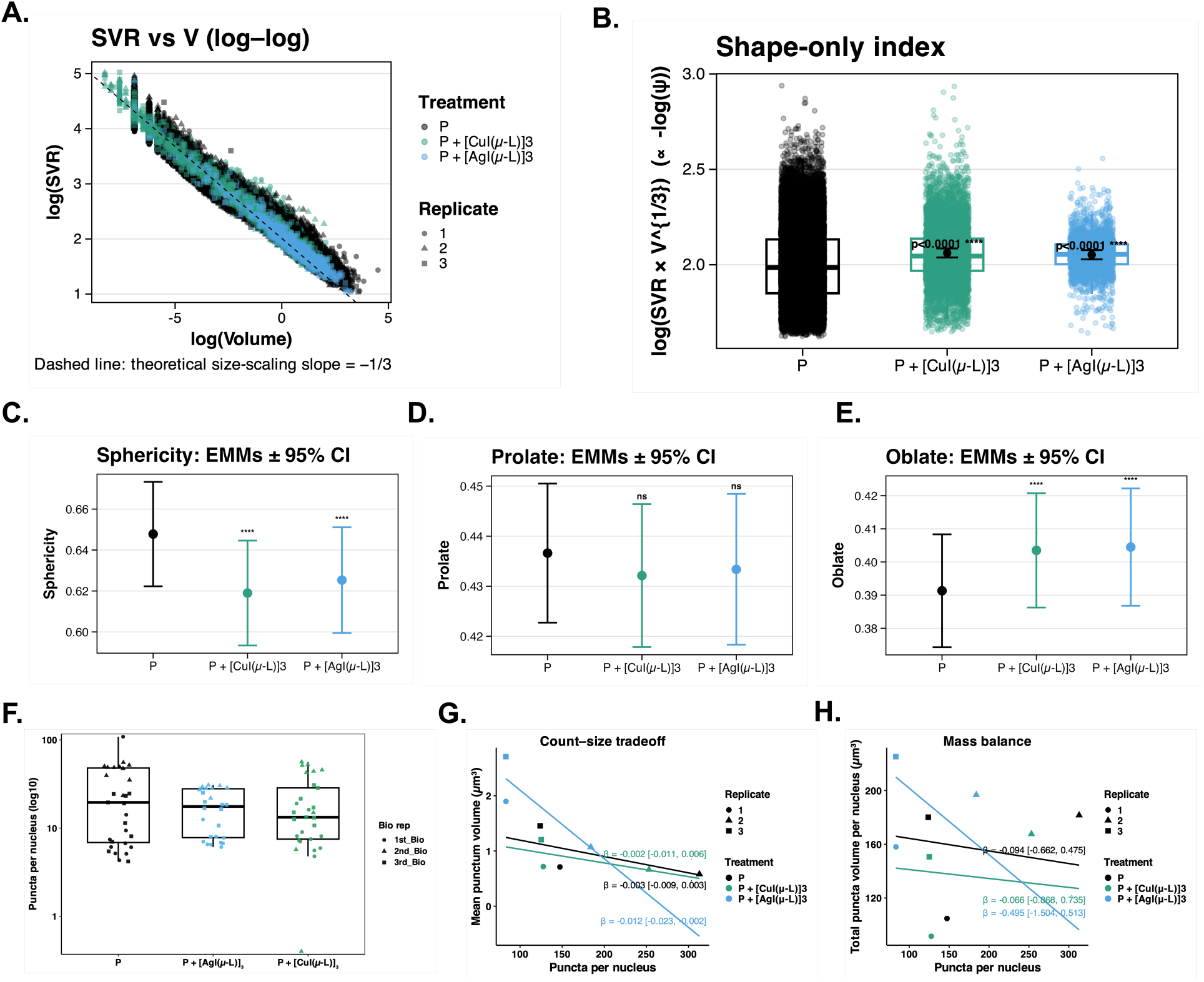
**[Ag^I^(**µ**-L)]_3_** results in less spherical, larger, more uniform-sizes puncta that show a trend towards growth in size over spread (n=3.) **(A-C)** Determination of size and shape. **(A)** log*(SVR*) vs log*(V*) graph shows the relationship between shape and size with a theoretical 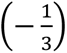 size-shape scale under geometrical similarity. At a common size, fitted lines representing **[Ag^I^(**µ**-L)]_3_** (blue) and **[Cu^I^(**µ**-L)]_3_** (teal) puncta show a vertical shift downwards as compared to control P (black) aggregates indicating smaller SVR values. This suggests aggregates are smoother/more compact. Treatment was significant F(2, 40,738) = 279.05, ANCOVA, p < 0.0001. **(B)** Shape only index with EMMs overlay. Graphing 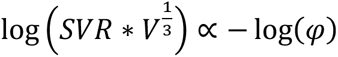 where *φ* = sphericity, **[Cu^I^(**µ**-L)]_3_** (teal) and **[Ag^I^(**µ**-L)]_3_** (blue) aggregates have larger values indicating lower sphericity. **(C)** Sphericity EMMs (mean [95% CI]) show: PFFs: 0.65 [0.62, 0.67]; **[Cu^I^(**µ**-L)]_3_** 0.62 [0.59, 0.64]; **[Ag^I^(**µ**-L)]_3_** 0.62 [0.60, 0.65] (****p < 0.0001). **(D)** Prolate. Values are unchanged. EMMs (mean [95% CI]) show PFFs: 0.44 [0.42, 0.45]; **[Cu^I^(**µ**-L)]_3_** shows a trending decrease 0.43 [0.42, 0.45] (p= 0.064); **[Ag^I^(**µ**-L)]_3_** shows no significant difference 0.43 [0.42, 0.45] (p=0.505) **(E)** Oblate. Values are increased for both treatments. EMMs (mean [95% CI]) show: PFFs: 0.39 [0.37, 0.41]; **[Cu^I^(**µ**-L)]_3_** 0.40 [0.39, 0.42]; **[Ag^I^(**µ**-L)]_3_** 0.40 [0.39, 0.42] (****p < 0.0001). **(F)** Puncta per nuclei. No overall significant difference is observed in the total puncta number (puncta per nucleus) (**[Cu^I^(**µ**-L)]_3_**, p = 0.48; **[Ag^I^(**µ**-L)]_3_**, p = 0.67 (Mixed-effects model, Dunnett pair-wise multiple comparisons.) **(G-H) [Ag^I^(**µ**-L)]_3_** increases puncta growth over puncta spread. **(G)** Count–size tradeoff. **[Ag^I^(**µ**-L)]_3_** reduced puncta per nucleus (IRR = 0.61, p<0.0001) but increased punctum size (GMR = 2.09, p<0.026) relative to P. **[Cu^I^(**µ**-L)]_3_** caused a modest reduction in puncta per nucleus (IRR = 0.89, p<0.011) without affecting size (GMR = 0.98, p=0.88.) **(H)** Mass balance. **[Ag^I^(**µ**-L)]_3_** does not affect the puncta volume per nucleus (TVNR = 1.29, p = 0.13), whereas **[Cu^I^(**µ**-L)]_3_** showed a small reduction (TVNR = 0.88, p<0.044).

We further investigated the significance of the SVR values in order to infer more information about aggregate characteristics. Here, we plotted a graph with log(*SVR*) on the y-axis and log(*V*)on the x-axis (Figure 8A.) This relationship is derived from the equation of SVR: 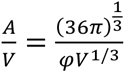 which yields 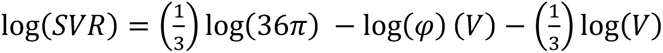, where *A* = surface area, *V* = volume, and *φ* = sphericity. According to this equation, a shape, such as a sphere, with a constant relationship between size and shape, would yield a graph with the equation: 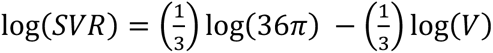 and therefore, a slope of 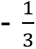. Higher values (larger log(*SVR*) values) would indicate that volume is larger than the surface area and therefore the shape of the proteins is more compact. Conversely, smaller SVR values would indicate that the surface area would be greater than the volume, leading to less compact aggregates. Similar observations were described for single peptide proteins characterized according to their SVR values.^69^

**Figure 8.**
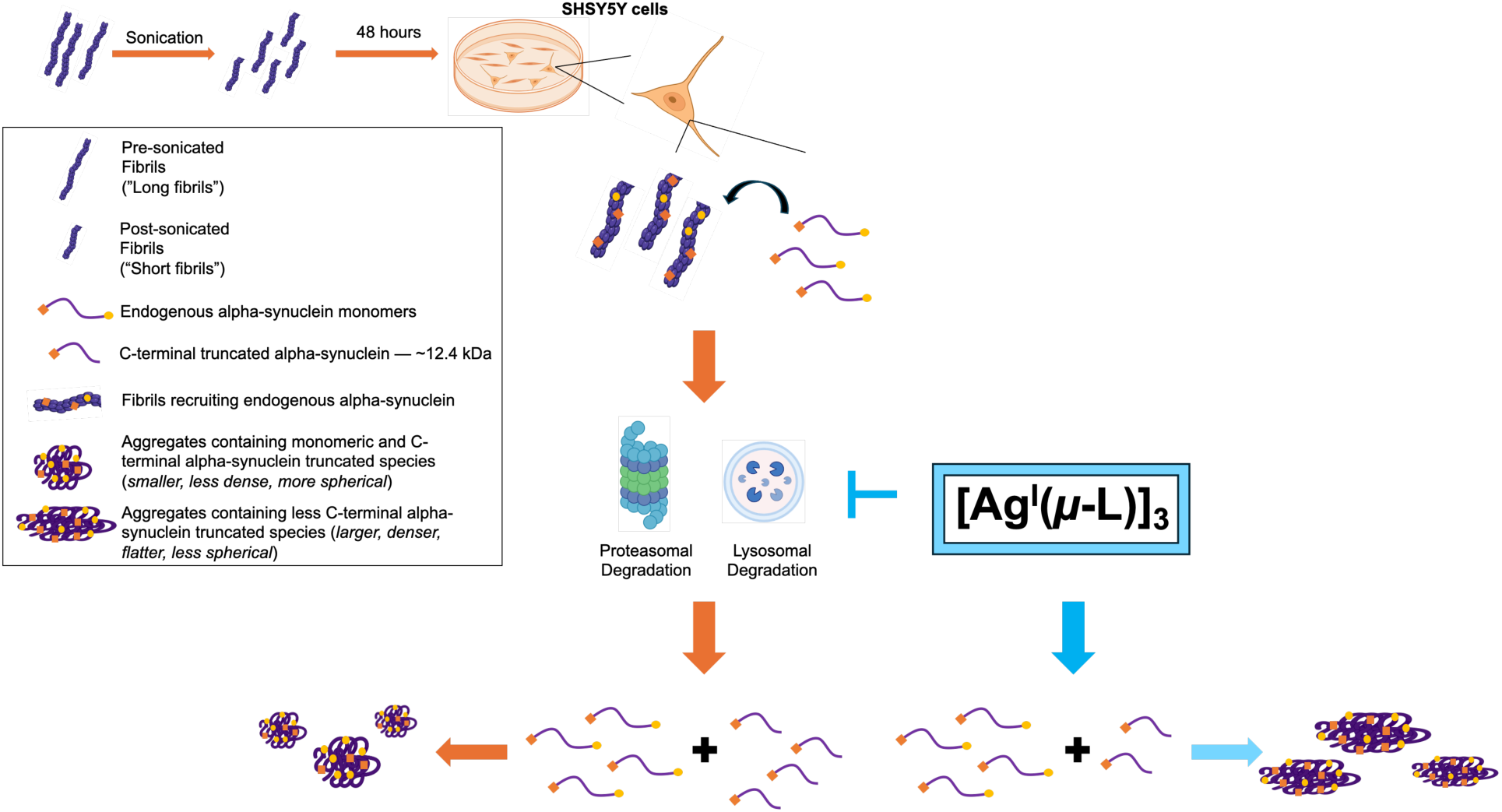
Hypothesis: **[Ag^I^(**µ**-L)]_3_** inhibits proteosomal or lysosomal degradation mechanisms, reducing the amount of the ∼12.4 kDa C-terminal alpha-synuclein truncation product, but increasing aggregate size (surface area, volume) and altering the shape leading to denser, flatter, and less spherical aggregates. Full length alpha-synuclein fibrils were initially sonicated to obtain shorter, more aggregate prone, alpha-synuclein fibrils. These fibrils accumulate endogenous alpha-synuclein (peptides containing yellow circles and orange squares) into larger aggregates over a 48-hour period that are degraded via proteasomal or lysosomal degradation mechanisms to produce C-terminally truncated alpha-synuclein (peptides containing only orange squares.) Our results suggest that **[Ag^I^(**µ**-L)]_3_** specifically reduces the ∼12.4 kDa protein species, suggesting that it targets proteasomal or lysosomal degradation proteins, such as calpain 1, caspase-1 or cathepsin B, L or D (illustrated with Biorender.)

Upon graphing the fitted lines for the **[Cu^I^(**µ**-L)]_3_** and **[Ag^I^(**µ**-L)]_3_** treatment groups (teal and blue, respectively), we saw distinct characteristics emerge at given sizes (log(*V*)). Here, **[Cu^I^(**µ**-L)]_3_** and **[Ag^I^(**µ**-L)]_3_** showed that for some aggregates, the y-axis values were shifted downwards when compared to PFFs only (black), indicating lower SVR values. Overlap for other aggregates made visualization difficult. We also observed that **[Ag^I^(**µ**-L)]_3_** showed clustering of aggregates towards larger log(*V*) values. In order to understand whether puncta may differ by shape, we plotted size-independent values (Figure 7B): 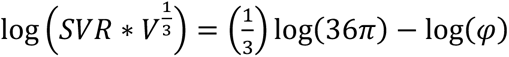. Here, larger *φ* (sphericity) values would result in smaller values on the y-axis. We found that PFFs (black) showed significantly smaller values as compared to both **[Cu^I^(**µ**-L)]_3_** and **[Ag^I^(**µ**-L)]_3_** (teal and blue, respectively). These results were confirmed in Figure 7C where average sphericity values showed that **[Cu^I^(**µ**-L)]_3_** and **[Ag^I^(**µ**-L)]_3_** have significantly smaller values (0.62) as compared to PFFs alone (0.65) (****p < 0.0001).

We also characterized aggregate shape by comparing prolate (elongated), oblate (disk-like), and sphericity (sphere-like) values. Interestingly, while **[Cu^I^(**µ**-L)]_3_** treatment resulted in a trending decrease for prolate values (0.43, p=0.064) as compared to PFFs alone (0.44), values were unchanged for **[Ag^I^(**µ**-L)]_3_** (0.43, p=0.505). Conversely, oblate values showed significant increases with both **[Cu^I^(**µ**-L)]_3_** (0.40) and **[Ag^I^(**µ**-L)]_3_** (0.40) as compared to PFFs alone (0.39) (****p < 0.0001). Together, these results suggest that sphericity may be lower for **[Ag^I^(**µ**-L)]_3_** treated aggregates which differ only according to oblate dimensions unlike **[Cu^I^(**µ**-L)]_3_** treatment which are perhaps more proportionally altered where changes occur in the oblate dimension as well as almost significantly altered in the prolate dimension. Together, this data suggests that **[Ag^I^(**µ**-L)]_3_** treatment yields more flattened, disk-like aggregate structures. We hypothesize this effect may be due to reduction in ∼12.4 kDa truncated species, where absence of this protein affects aggregation chemistry.

As our results have shown that **[Ag^I^(**µ**-L)]_3_** treatment leads to increased aggregate size, we next investigated whether aggregate numbers differed between treatments and the effect on aggregate spread. We saw that although the average number of puncta/nuclei were slightly smaller for **[Cu^I^(**µ**-L)]_3_** and **[Ag^I^(**µ**-L)]_3_** treated cells, there was no significant difference in these average values as compared to PFFs alone (Figure 7F) (p=0.48 and p=0.67, respectively.) However, given that both treatments result in larger aggregates, we aimed to better understand the relationship between individual aggregate size and puncta/nuclei (“puncta spread”). We found that given a smaller number of puncta/nuclei (IRR = 0.61, p<0.0001), the average size of **[Ag^I^(**µ**-L)]_3_** aggregates trended towards greater values (GMR =2.09, p<0.026) while **[Cu^I^(**µ**-L)]_3_** caused a modest reduction in puncta/nucleus (IRR = 0.89, p<0.011) without affecting size (GMR = 0.98, p=0.88.) The cumulative aggregate volume showed that **[Ag^I^(**µ**-L)]_3_** did not affect the total puncta mass volume (TVNR = 1.29, p = 0.13), whereas **[Cu^I^(**µ**-L)]_3_** showed a small reduction (TVNR = 0.88, p<0.044) (Figure 7H, Table 1.) This suggests that while **[Ag^I^(**µ**-L)]_3_** leads to aggregate growth, it does not increase the total amount of aggregates formed.

Altogether, we see that **[Ag^I^(**µ**-L)]_3_** treatment leads to reduction of the ∼12.4 kDa C-terminal truncated species and results in larger, more uniform, less spherical, more oblate aggregates, which appear to favor growth over the spread of aggregation. Given that C-terminal truncated species impact alpha-synuclein aggregation by inducing aggregation and affecting fibrillic structure/shape,^35,36^ our data further suggests that changing the dynamics of monomer to ∼12.4 kDa C-terminally truncated species affects PFFs-induced alpha-synuclein protein aggregation.

## Discussion

Alpha-synuclein pre-formed fibrils (PFFs) are a widely used model for the study of Parkinson’s related protein aggregation.^18,19^ Treatment with PFFs in both cell and mouse models has been shown to lead to propagation of aggregation, including spread from the gut to the brain, as well as induce insoluble protein aggregate formation.^18,19,23,56^ Recent findings have shown that alpha-synuclein truncations at the amino (N-) or carboxy (C-) terminus can determine alpha-synuclein’s aggregation propensity, with C-terminal truncations contributing strongly towards increased aggregation.^35,37,40^

As alpha-synuclein protein aggregation is one of the characteristics of Parkinson’s biochemistry, research has focused on methods to reduce aggregation, such as removal of C-terminally truncated species. Previous work found that the Anle138b (**HL**) pyrazole chemical interacts with toxic alpha-synuclein oligomeric species as well as fibrils, and reduces alpha-synuclein protein aggregation.^59^ In order to determine if metal chelation causes alternate effects on aggregation, we have synthesized the **HL** analogues: **H_2_LClO_4_**, **[Cu^I^(**µ**-L)]_3_**, and **[Ag^I^(**µ**-L)]_3_**. We then treated SHSY5Y human dopaminergic cells with these compounds to determine their bioviability and effect on alpha-synuclein protein aggregation.

We first assessed toxicity with the MTT assay and asked whether the compounds protect cells from a moderately toxic PFFs dose (15 µg/mL, or ∼1 µM) (Figure 3, S11). Given that cells were treated concomitantly with chemical compounds and PFFs, we hypothesized this would allow for protein-chemical interactions to occur as previously observed between Anle138b and oligomers or fibrils^59,70^ leading to a potential interaction that would reduce PFF induced toxic effects. While we did observe that at certain concentrations for **HL** and **H_2_LClO_4_** (50 µM) and **[Cu^I^(**µ**-L)]_3_** (25 uM), addition of PFFs resulted in slightly higher MTT values than with compound alone, these results did not achieve a significant effect (Figure 3.) We did not observe similar effects for **[Ag^I^(**µ**-L)]_3_**, indicating that other mechanisms may be involved.

Instead, we observed that **[Ag^I^(**µ**-L)]_3_** had the strongest and only effect of reducing a ∼12.4 kDa alpha-synuclein protein species. Through treatment with two separate antibodies — one recognizing residues 15-123 kDa (BD antibody) and the other recognizing residues only within the C-terminus (surrounding proline 129, and including residues 122 and 123) (Figure 5 A, C, and E) — and application of the Edman Sequencing method (Figure S13), found that the ∼12.4 kDa species is C- and not N-terminally truncated. Importantly, differences between alpha-synuclein antibody detection patterns were previously observed by others as well, and it has been indicated that various post-translational modifications, including C-terminal truncations, can affect alpha-synuclein detection.^58^

Of note is also the abundance of ∼15 kDa band in cells treated with PFFs (Figure 5 and 6, blue arrow). This band appears to be the most concentrated when compared to all other protein bands in the lane. Because monomeric alpha-synuclein is usually measured higher at 19 kDa (17 kDa in our blots), we hypothesize that the 15 kDa band may also be a truncation product, isoform, or an otherwise post-translationally modified alpha-synuclein species.^71^ If this is indeed a truncated protein, the detection by both antibodies signifies cleavage would likely occur outside the recognizable immunogen for either antibody. Alternatively, the prominent accumulation of the ∼15 kDa may suggest that lysosomal or proteasomal homeostasis is altered as previously reported in the context of PD.^40,72^

Mechanisms altering protein degradation pathways have been shown to decrease alpha-synuclein C-terminal truncation. A previous report showed that caspase-1 inhibition reduced a 12 kDa C-terminally truncated alpha-synuclein species and consequentially, decreased higher molecular weight aggregates in a mouse model of multiple system atrophy (MSA).^34^ Additional literature suggests that truncation resulting in a ∼12.4 kDa protein could be due to cleavage at several sites, including: 1) 122/123 by calpain 1, caspase 1, asparagine endopeptidase, or cathepsins B or L; 2) 124/125 by cathepsin D; or 3) 125/126 by cathepsin L.^40^ However, unlike these reports, including one showing that reducing metalloprotease cleavage activity results in a reduction of alpha-synuclein aggregation,^73^ our data suggests that reduction of the ∼12.4 kDa species by **[Ag^I^(**µ**-L)]_3_** actually yields enlarged aggregates with great surface area and volume (Figure 7 A,B.)

Together, this suggests a more nuanced picture. While **[Ag^I^(**µ**-L)]_3_** may decrease mechanisms related to protein degradation, causing a decrease in ∼12.4 kDa species, it may also affect protein dynamics in such a way as to modify protein size when the species conglomerate to form aggregates. Previously, it was shown that truncated alpha-synuclein species (1-108) form fibrils with tighter β-sheet structures than those formed from full length monomers.^35^ This suggests that the aggregate structure depends largely upon how the monomers or truncated alpha-synuclein proteins interact with one another. Thus, in a mixed population of protein species, interactions taking place between monomers and C-terminally truncated species could result in aggregates of different structure and shape apart from aggregates formed from a population of species that contain less truncated species. We thus see from our results that the surface to volume ratio (SVR) is significantly smaller with **[Ag^I^(**µ**-L)]_3_** treatment (Figure 7C), suggesting that proteins interact differently to form more compact aggregates. Furthermore, the more oblate but lower spherical character of **[Ag^I^(**µ**-L)]_3_** treated proteins suggests that a directionality of growth could be present, resulting in flatter/disk-like aggregates. Finally, we see that aggregate growth is preferred over aggregate spread in a significantly trending manner (Figure 7 F-E) suggesting that **[Ag^I^(**µ**-L)]_3_** may slow aggregate seeding by reducing the ∼12.4 kDa C-terminal truncated species.

Our work therefore highlights that complexation of Anle138b with silver causes an outcome unique from the effect of the original Anle138b compound. Unlike previously published reports which demonstrate that reduction of C-terminally truncated alpha-synuclein species leads to decreased aggregation, **[Ag^I^(**µ**-L)]_3_** treatment causes aggregates to enlarge. We hypothesize that **[Ag^I^(**µ**-L)]_3_** inhibits alpha-synuclein cleavage through action on protein degradation mechanisms leading to reduction of the ∼12.4 kDa C-terminally truncated species (Figure 8). Our results further suggest that treatment of cells with PFFs and **[Ag^I^(**µ**-L)]_3_** concomitantly results in increased alpha-synuclein aggregate size, differential shape, and a preference for aggregate growth over spread. Future work will explore effects on protein degradation pathways and alternative experimental methods to shed greater light on these results.

## Conclusions

In this paper, we report our findings that coordination of **HL** to silver results in a novel compound, **[Ag^I^(**µ**-L)]_3_**. Unlike **HL** and the other synthesized compounds characterized here, **[Ag^I^(**µ**-L)]_3_** treatment of SHSY5Y cells together with alpha-synuclein PFFs resulted in a unique effect: Unlike **HL**, **[Ag^I^(**µ**-L)]_3_** leads to a reduction of a C-terminally truncated alpha-synuclein protein species of approximately 12.4 kDa. While the scope of this paper focused on the characterization and the bioviability of the synthesized compounds and the alpha-synuclein aggregates, future work with aim to explore the biological significance of the enlarged alpha-synuclein aggregates as well as explore **[Ag^I^(**µ**-L)]_3_** chemical properties, such as luminescence, which will allow *in vitro* monitoring of aggregation. We also hypothesize that our results may depend upon the nature of our experimental model i.e. concomitant treatment of compound and PFFs versus protein overexpression in transgenic animals, as previously reported.^34^ Overall, we demonstrate that **[Ag^I^(**µ**-L)]_3_** increases aggregate size despite reducing a C-terminal truncation species. Further research into chemicals which exert novel effects upon truncation species and consequentially, alpha-synuclein aggregation, may prove important for uncovering a greater understanding of the models used for Parkinson’s research. In this way, we propose that **[Ag^I^(**µ**-L)]_3_** can uniquely contribute toward this field.

## Acknowledgements

KLR and SH were supported by a U.S. Nuclear Regulatory Commission (NRC) fellowship grant No. 31310018M0012 awarded to FIU.

Cell bioviability and protein experiments were supported by the NIH NIA T32 AG000266 Grant. Edman sequencing was performed by The Protein Facility of the Iowa State University Office of Biotechnology.

## References

(1) Zafar, S.; Yaddanapudi, S. Parkinson Disease. In StatPearls; 2023. https://www.ncbi.nlm.nih.gov/books/NBK470193/ (accessed 2025-07-21).

(2) Dorsey, E. R.; Sherer, T.; Okun, M. S.; Bloem, B. R. The Emerging Evidence of the Parkinson Pandemic Brundin P, Langston JW, Bloem BR, editors. J. Parkinsons Dis. 2018, 8, S3–S8. 10.3233/JPD-181474.

(3) Fan, H.; Sheng, S.; Zhang, F. New Hope for Parkinson’s Disease Treatment: Targeting Gut Microbiota. CNS Neurosci. Ther. 2022, 28, 1675–1688. 10.1111/cns.13916.

(4) Maeda, T.; Nagata, K.; Satoh, Y.; Yamazaki, T.; Takano, D. High Prevalence of Gastroesophageal Reflux Disease in Parkinson’s Disease: A Questionnaire-Based Study. J. Parkinson’s Dis. 2013, 742128 (6 pages). 10.1155/2013/742128.

(5) Bock, M. A.; Brown, E. G.;Zhang, L.; Tanner, C. Association of Motor and nonmotor Symptoms With Health-Related Quality of Life in a Large Online Cohort of People With Parkinson Disease. Neurol. 2022, 98, e2194–e3303. 10.1212/WNL.0000000000200113.

(6) Pereira, F. C.; Ge, X.; Kristensen, J. M.; Kirkegaard, R. H.; Maritsch, K.; Szamosvári, D.; Imminger, S.; Seki, D.; Shazzad, J. B.; Zhu, Y.; Decorte, M.; Hausmann, B.; Berry, D.; Wasmund, K.; Schintlmeister, A.; Böttcher, T.; Cheng, J.-X.; Wagner, M. The Parkinson’s disease drug entacapone disrupts gut microbiome homoeostasis via iron sequestration. Nature Microbiol. 2024, 9, 3165–3183. 10.1038/s41564-024-01853-0.

(7) Braak, H.; Rüb, U.; Gai, W. P.; Del Tredici, K. Idiopathic Parkinson’s Disease: Possible Routes by Which Vulnerable Neuronal Types May Be Subject to Neuroinvasion by an Unknown Pathogen. J. Neural Transm. 2003, 110, 517–536. 10.1007/s00702-002-0808-2.

(8) Svensson, E.; Horvath, L.; Åsberg Johnels, J.; Goulding, N.; Kwon, B.-J.; Holmqvist, S.; Mulkidzanyan, S.; Richter, R. P.; Lindhagen-Persson, M.; Nordström, E.;, et al. Vagotomy and Subsequent Risk of Parkinson’s Disease: Vagotomy and Risk of PD. Ann. Neurol. 2015, 78, 522–529. 10.1002/ana.24448.

(9) Del Tredici, K.; Braak, H. Review: Sporadic Parkinson’s Disease: Development and Distribution of α-Synuclein Pathology. Neuropathol. Appl. Neurobiol. 2016, 42, 33–50. 10.1111/nan.12298.

(10) Van Den Berge, N.; Ferreira, N.; Phillips, R. J.; Tjaden, A.; Meyer, C.; Sidhu, M.; Hesta, L.; Libby, R. T.; Nuydens, R.; Vandenberghe, R.;, et al. Evidence for Bidirectional and Trans-Synaptic Parasympathetic and Sympathetic Propagation of Alpha-Synuclein in Rats. Acta Neuropathol. 2019, 138, 535–550. 10.1007/s00401-019-02040-w.

(11) Horsager, J.; Hansen, D.; Peters, N.; Sommerauer, M.; Nußbaum, S.; Hertz, J. M.; Holm, I. E.; Serradell, M.; Møller, C.; Hinz, R.;, et al. Brain-First Versus Body-First Parkinson’s Disease: A Multimodal Imaging Case-Control Study. Brain 2020, 143, 3077–3088. 10.1093/brain/awaa238.

(12) Lai, Y.; Kim, J.; Chung, K. C.; Kishimoto, S.; Kim, D.; Lim, D. H.; Kang, D.; Lee, S. K.; Han, H.; Han, K. H.;, et al. Nonaggregated α-Synuclein Influences SNARE-Dependent Vesicle Docking via Membrane Binding. Biochemistry 2014, 53, 3889–3896. 10.1021/bi5002536.

(13) Chandra, R.; Samuelson, I.; Zhang, M.; Szimmtenings, P.; Schwendeman, A.; Kordower, J. H.; Sortwell, C. E.; Lee, J. C. α-Synuclein in Gut Endocrine Cells and Its Implications for Parkinson’s Disease. JCI Insight 2017, 2, e92295. 10.1172/jci.insight.92295.

(14) Man, W. K.; Yu, S.; Wang, X.; Neng, L.; Cheng, H.; Wang, F.; Wang, P.; Chen, C. M.; Liang, S.; Li, J.;, et al. The Docking of Synaptic Vesicles on the Presynaptic Membrane Induced by α-Synuclein Is Modulated by Lipid Composition. Nat. Commun. 2021, 12, 927. 10.1038/s41467-021-21027-4.

(15) Stephens, A. D.; O’Neill, M. J.; Foden, H. R.; Smedley, K. C.; Kye, C.; Hirst, W. D.; Volles, M. P. α-Synuclein Fibril and Synaptic Vesicle Interactions Lead to Vesicle Destruction and Increased Lipid-Associated Fibril Uptake into iPSC-Derived Neurons. Commun. Biol. 2023, 6, 526. 10.1038/s42003-023-04884-1.

(16) Cuervo, A. M.; Stefanis, L.; Fredenburg, R.; Lansbury, P. T.; Sulzer, D. Impaired Degradation of Mutant α-Synuclein by Chaperone-Mediated Autophagy. Science 2004, 305, 1292–1295. 10.1126/science.1101738.

(17) Martinez-Vicente, M.; Talloczy, Z.; Kaushik, S.; de Vries, R.; Arias, E.; Lawrence, R. A.; Paine, A. L.; Cullen, A.; S Fraser, K.; Wang, F.;, et al. Dopamine-Modified α-Synuclein Blocks Chaperone-Mediated Autophagy. J. Clin. Invest. 2008, 118, 772–782. 10.1172/JCI32806.

(18) Duffy, M. F.; Collier, T. J.; Patterson, J. R.; Stoll, A. D.; Kemp, C. J.; Allen Reish, H. E.; Madaj, Z.; Meshul, C. K.; Sortwell, C. E.; Standaert, D. G.;, et al. Quality over Quantity: Advantages of Using Alpha-Synuclein Preformed Fibril Triggered Synucleinopathy to Model Idiopathic Parkinson’s Disease. Front. Neurosci. 2018, 12, 621. 10.3389/fnins.2018.00621.

(19) Uemura, N.; Peelaerts, W.; Larkin, M. J.; Luk, K. C.; Jude, F.; Ste Historical, E.; Dufour, A.; Van der Perren, A.; Baekelandt, V.; Taylor, J. P.;, et al. Inoculation of α-Synuclein Preformed Fibrils into the Mouse Gastrointestinal Tract Induces Lewy Body-Like Aggregates in the Brainstem via the Vagus Nerve. Mol. Neurodegener. 2018, 13, 21. 10.1186/s13024-018-0257-5.

(20) Kim, B.-J.; Noh, H.-R.; Jeon, H.; Park, S.-M. Monitoring a-synnuclein Aggregation Induced by Preformed a-synuclein Fibrils in an *In Vitro* Model System. Exp. Neurobiol. 2023, 32, 147–156. 10.5607/en23007

(21) Horsager, J.; Parkinsonism 2024

(22) Alam, P.; Bousset, L.; Melki, R.; Otzen, D. E. α-Synuclein Oligomers and Fibrils: A Spectrum of Species, a Spectrum of Toxicities. J. Neurochem. 2019, 150, 522–534. 10.1111/jnc.14808.

(23) Cascella, R.; Campana, M.; Cacace, M.; De Simone, M.; Pagano, B.; Arcucci, M. V.; Di Martino, S.; D’Onofrio, G.; Palamà, I.; Roviello, S.;, et al. The Release of Toxic Oligomers from α-Synuclein Fibrils Induces Dysfunction in Neuronal Cells. Nat. Commun. 2021, 12, 1814. 10.1038/s41467-021-21937-3.

(24) Choi, M. L.; Kim, J. S.; Gwon, Y.; Kim, J.; Jang, S.; Jeong, J.; Jeong, H.; Lee, H.-J.; Park, Y. S.; Park, Y.;, et al. Pathological Structural Conversion of α-Synuclein at the Mitochondria Induces Neuronal Toxicity. Nat. Neurosci. 2022, 25, 1134–1148. 10.1038/s41593-022-01140-3.

(25) Allen Reish, H. E.; Standaert, D. G. Role of α-Synuclein in Inducing Innate and Adaptive Immunity in Parkinson Disease. J. Parkinsons Dis. 2015, 5, 1–19. 10.3233/JPD-140491.

(26) Labrie, V.; Brundin, P. Alpha-Synuclein to the Rescue: Immune Cell Recruitment by Alpha-Synuclein During Gastrointestinal Infection. J. Innate Immun. 2017, 9, 437–440. 10.1159/000479653.

(27) Barbut, D.; Stolzenberg, E.; Zasloff, M. Gastrointestinal Immunity and Alpha-Synuclein van Laar T, editor. J. Parkinsons Dis. 2020, 9, S313–S322. 10.3233/JPD-191702.

(28) Yamin, G.; Uversky, V. N.; Fink, A. L. Nitration Inhibits Fibrillation of Human α-Synuclein in Vitro by Formation of Soluble Oligomers. FEBS Lett. 2003, 542, 147–152. 10.1016/S0014-5793(03)00367-3.

(29) Wördehoff, M. M.; Geers, H.; Scheidt, H. A.; Finke, J. M.; Schierhorn, A.; Hause, G.; Stichel, J.; Smits, R.; Schlenzig, D.; Göricke, B.;, et al. Opposed Effects of Dityrosine Formation in Soluble and Aggregated α-Synuclein on Fibril Growth. J. Mol. Biol. 2017, 429, 3018–3030. 10.1016/j.jmb.2017.09.005.

(30) Al-Hilaly, Y. K.; King, Z. A.; Linton, P.; Bleiholder, R.; Ganuelas, M. L.; Khalaf, A. I.; Parkinson, G. N.; Tuttle, M. D.; Sovago, M.; Jakes, R.;, et al. The Involvement of Dityrosine Crosslinking in α-Synuclein Assembly and Deposition in Lewy Bodies in Parkinson’s Disease. Sci. Rep. 2016, 6, 39171. 10.1038/srep39171.

(31) Shahmoradian, S. H.; Genoud, C.; Quattrini, A.; Jiménez-Ferrer, I.; Rodríguez-Andrés, C.; Castano-Díez, D.; Albert, S.; Bousset, L.; Carpentier, G.; Pieri, L.;, et al. Lewy Pathology in Parkinson’s Disease Consists of Crowded Organelles and Lipid Membranes. Nat. Neurosci. 2019, 22, 1099–1109. 10.1038/s41593-019-0423-2.

(32) Siderowf, A.; Concha-Marambio, L.; Lafontant, D.-E.; Fagerqvist, T.; Hart, E.; Freedman, S. N.; Simms, E.; Budd Haeberlein, S.; Dujardin, S.; Espay, A. J.;, et al. Assessment of Heterogeneity Among Participants in the Parkinson’s Progression Markers Initiative Cohort Using α-Synuclein Seed Amplification: A Cross-Sectional Study. Lancet Neurol. 2023, 22, 407–417. 10.1016/S1474-4422(23)00109-6.

(33) Yan, F. Gastrointestinal Nervous System α-Synuclein as a Potential Biomarker of Parkinson Disease. Medicine, 2018, 97, e11337. 10.1097/MD.0000000000011337

(34) Bassil, F.; Guerin, P.; Dufour, N.; Conde, S.; Rousseau, D.; Dehay, B.; Li, J. Y.; Bezard, E.; Przedborski, S.; Olanow, C. W.;, et al. Reducing C-Terminal Truncation Mitigates Synucleinopathy and Neurodegeneration in a Transgenic Model of Multiple System Atrophy. Proc. Natl. Acad. Sci. 2016, 113, 9593–9598. 10.1073/pnas.1609291113.

(35) Iyer, A.; Leong, S. L.; Ng, L. P.; Lee, Y.-H.; Lim, L. K.; Liu, K.; Wenk, M. R.; Lim, K. H. C-Terminal Truncated α-Synuclein Fibrils Contain Strongly Twisted β-Sheets. J. Am. Chem. Soc. 2017, 139, 15392–15400. 10.1021/jacs.7b07403.

(36) Ma, L.; Huang, B.; Yuan, Z.; Huang, H.; Wang, X. C-Terminal Truncation Exacerbates the Aggregation and Cytotoxicity of α-Synuclein: A Vicious Cycle in Parkinson’s Disease. Biochim. Biophys. Acta BBA - Mol. Basis Dis. 2018, 1864, 3714–3725. 10.1016/j.bbadis.2018.10.003.

(37) Sorrentino, Z. A.; Xia, Y.; Ko, H. S.; Zhang, L.; Bleiholder, R.; Dunker, A. K.; Canty, A. J.; Lee, J. C.; Lee, V. M. Y.; Giasson, B. I. Physiological C-Terminal Truncation of α-Synuclein Potentiates the Prion-Like Formation of Pathological Inclusions. J. Biol. Chem. 2018, 293, 18914–18932. 10.1074/jbc.RA118.005603.

(38) Farzadfard, A.; Ghantasala, K. S.; Rospigliosi, C. C.; Roy, D.; Jha, N.; Subramanian, S.; Mishra, V. S.; Varthya, V.; Das, S.; S, L. V.;, et al. The C-Terminal Tail of α-Synuclein Protects Against Aggregate Replication but Is Critical for Oligomerization. Commun. Biol. 2022, 5, 123. 10.1038/s42003-022-03059-8.

(39) Zhang, C.; Lu, M.; Ma, L.; Huang, H.; Wang, X. C-Terminal Truncation Modulates α-Synuclein’s Cytotoxicity and Aggregation by Promoting the Interactions With Membrane and Chaperone. Commun. Biol. 2022, 5, 798. 10.1038/s42003-022-03768-0.

(40) Sorrentino, Z. A.; Giasson, B. I. The Emerging Role of α-Synuclein Truncation in Aggregation and Disease. J. Biol. Chem. 2020, 295, 10224–10244. 10.1074/jbc.REV120.011743.

(41) Games, D.; Lee, H. G.; Adams, S.; Barbour, R.; Bartholow, K.; Freedman, S.; Galimi, F.; Hutter, J.; Keane, A.; Liu, H.;, et al. Reducing C-Terminal-Truncated Alpha-Synuclein by Immunotherapy Attenuates Neurodegeneration and Propagation in Parkinson’s Disease-Like Models. J. Neurosci. 2014, 34, 9441–9454. 10.1523/JNEUROSCI.5314-13.2014.

(42) Mittur, A.; Gupta, S.; Modi, N. B. Pharmacokinetics of Rytary®, an Extended-Release Capsule Formulation of Carbidopa–Levodopa. Clin. Pharmacokinet. 2017, 56, 999–1014. 10.1007/s40262-017-0511-y.

(43) Levin, J.; Giese, A.; Respondek, G.; Giese, R.; Stautner, S.; Zeuner, K.; Oertel, W. H.; Mansmann, U.; Klopstock, T.; Trenkwalder, C.;, et al. Safety, Tolerability and Pharmacokinetics of the Oligomer Modulator Anle138b With Exposure Levels Sufficient for Therapeutic Efficacy in a Murine Parkinson Model: A Randomised, Double-Blind, Placebo-Controlled Phase 1a Trial. EBioMedicine 2022, 80, 104021. 10.1016/j.ebiom.2022.104021.

(44) Levin, J.; Giese, A.; Respondek, G.; Giese, R.; Stautner, S.; Zeuner, K.; Oertel, W. H.; Mansmann, U.; Klopstock, T.; Trenkwalder, C.;, et al. Anle138b-P1-02: A Randomised, Double-Blinded, Placebo-Controlled Phase 1b Study to Investigate Safety, Tolerability, Pharmacokinetics and Pharmacodynamics of the Oligomer Modulator Anle138b in Parkinsońs Disease. Mov. Disord. 2023, 38 (suppl 1). https://www.mdsabstracts.org/abstract/anle138b-p1-02-a-randomised-double-blinded-placebo-controlled-phase-1b-study-to-investigate-safety-tolerability-pharmacokinetics-and-pharmacodynamics-of-the-oligomer-modulator-anle138b-in-parkins/ (accessed 2025-01-06).

(45) Levin, J.; Giese, A.; Respondek, G.; Giese, R.; Stautner, S.; Zeuner, K.; Oertel, W. H.; Mansmann, U.; Klopstock, T.; Trenkwalder, C.;, et al. Targeting Oligomer Pathology of Alpha-Synuclein – A Study Evaluating the Safety and Efficacy of Emrusolmin in Patients With Multiple System Atrophy. Mov. Disord. 2024, 39 (suppl 1). https://www.mdsabstracts.org/abstract/topas-msa-targeting-oligomer-pathology-of-alpha-synuclein-a-study-evaluating-the-safety-and-efficacy-of-emrusolmin-in-patients-with-multiple-system-atrophy/ (accessed 2025-01-06).

(46) Deeg, A. A.; Reiner, A. M.; Schmidt, F.; Schueder, F.; Ryazanov, S.; Ruf, V. C.; Giller, K.; Becker, S.; Leonov, A.; Griesinger, C.;, et al. Anle138b and Related Compounds Are Aggregation Specific Fluorescence Markers and Reveal High Affinity Binding to α-Synuclein Aggregates. Biochim. Biophys. Acta BBA - Gen. Subj. 2015, 1850, 1884–1890. 10.1016/j.bbagen.2015.05.021.

(47) Antonschmidt, L.; Rönnefahrt, N.; Giller, K.; Sticht, H.; Griesinger, C.; Becker, S.; Zweckstetter, M. The Clinical Drug Candidate Anle138b Binds in a Cavity of Lipidic α-Synuclein Fibrils. Nat. Commun. 2022, 13, 5385. 10.1038/s41467-022-32797-w.

(48) Dervişoğlu, R.; Giller, K.; Griesinger, C.; Becker, S.; Leonov, A.; Zweckstetter, M. Anle138b Interaction in α-Synuclein Aggregates by Dynamic Nuclear Polarization NMR. Methods 2023, 214, 18–27. 10.1016/j.ymeth.2023.04.002.

(49) Lopes, L. G. F.; Carvalho, E. M.; Sousa, E. H. S. A bioinorganic chemistry perspective on the roles of metals as drugs and targets against *Mycobacterium Tuberculosis* – a journey of opportunities. Dalton Trans. 2020, 49, 15988. 10.1-39/d0dt01365j.

(50) Sierra, M. A.; Csarrubios, L.; de la Torre, M. C. Bio-Organometallic Derivatives of Antibacterial Drugs. Chem. Eur. J. 2019, 25, 7232–7242. 10.1002/chem.201805985.

(51) Riccardi, L.; Genna, V.; De Vivo, M. Metal-ligand interactions in drug design. Nature Rev. 2018, 2, 100–112. 10.1038/s41570-018-0018-6

(52) Renfrew, A. K. Transition metal complexes with bioactive ligands: mechanisms for selective ligand release and applications for drug delivery. Metallomics, 2014, 6, 1324–1335. 10.1039/c4mt00069b.

(53) Roecker, A. J.; Schirripa, K. M.; Loughran, H. M.; Tong, L.; Liang, T.; Fillgrove, K. E.; Kuo, Y.; Bleasby, K.; Collier, H.; Altman, M. D.; Ford, M. C.; Drolet, R. E.; Cosden, M.; Jinn, S.; Hatcher, N. G.; Yao, L.; Kandebo, M.; Vardigan, J. D.; Flick, R. B.; Liu, X.; Minnick, C.; Price, L. A.; Watt, M. L.; Lemaire, W. Burlein, C.; Adam, G. C.; Austin, L. A.; Marcus, J. N.; Smith, S. M. Fraley, M. E. Pyrazole Ureas as Low Dose, CNS Penetrant Glucosylceramide Synthase Inhibitors for the Treatment of Parkinsin;s Disease. ACS Med. Chem. Lett. 2023, 14, 146–155. 10.1021/acsmedchemlett.2c00441.

(54) Lésniak, R. K.; Nichols, R. J.; Schonemann, M.; Zhao, J.; Gajera, C. R.; Lam, G.; Nguyen, K. C.; Langston, J. W.; Smith, M.; Montine, T. J. Discovery of 1H-Pyrazole Biaryl Sulfonamides as Novel G2019S-LKKK2 Kinase Inhibitors. ACS Med. Chem. Lett. 2022, 13, 981–988. 10.1021/acsmedchemlett.2c00116.

(55) Candito, D. A.; Simov, V.; Gulati, A.; Kattar, S.; Chau, R. W.; Lapointe, B. T.; Methot, J. L.; DeMong, D. E.; Graham, T. H.; Kurukulasuriya, R.; Keylor, M. H.; Tong, L.; Moriello, G. J.; Acton, J. J.; Pio, B.; Liu, W.; Scott, J. D.; Ardolino, M. J.; Martinot, T. A.; Maddess, M. L; Yan, X.; Gunaydin, H.; Palte, R. L.; McMinn, S. E.; Nogle, L.; Yu, H.; Minnihan, E. C.; Lesburg, C. A.; Liu, P.; Su, J.; Hedge, L. G.; Moy, L. Y.; Woodhouse, J. D.; Faltus, C. A.; Xiong, T.; Ciaccio, P.; Piesvaux, J. A.; Otte, K. M.; Kennedy, M. E.; Bennett, D. J.; DiMauro, E. F.; Fell, M. J.; Neelamkavil, S.; Wood, H. B.; Fuller, P. H.; Ellis, J. M. Discovery and Optimization of Potent, Selective, and Brain-Penetrant 1-Heteroaryl-1H-Indazole LRRK2 Kinase Inhibitors for the Treatment of Parkinson’s Disease. J. Med. Chem. 2022, 65, 16801–16817. ACS Med. Chem. Lett. 2022, 13, 981-988. 10.1021/acs.jmedchem.2c01605.

(56) Kim, B. J.; Noh, H. R.; Jeon, H.; Park, S. M. Monitoring α-Synuclein Aggregation Induced by Preformed α-Synuclein Fibrils in an In Vitro Model System. Exp. Neurobiol. 2023, 32, 147–156. 10.5607/en23007.

(57) Bisaglia, M.; Bubacco, L. Copper Ions and Parkinson’s Disease: Why Is Homeostasis So Relevant? Biomolecules 2020, 10, 195. 10.3390/biom10020195.

(58) Altay, M. F.; Fagerqvist, T.; Concha-Marambio, L.; Tafsir, A.; Ingelsson, M.; Hyman, B. T.; St. George-Hyslop, P. H.; Hirst, W. D.; Dujardin, S.; Budd Haeberlein, S.;, et al. Development and Validation of an Expanded Antibody Toolset That Captures Alpha-Synuclein Pathological Diversity in Lewy Body Diseases. Neuroscience 2022. http://biorxiv.org/lookup/doi/10.1101/2022.05.26.493598 (accessed 2025-07-21).

(59) Wagner, J.; Ryazanov, S.; Leonov, A.; Levin, J.; Shi, S.; Schmidt, F.; Schmid, B.; Griesinger, C.; Giese, A.; Zweckstetter, M. Anle138b: A Novel Oligomer Modulator for Disease-Modifying Therapy of Neurodegenerative Diseases Such as Prion and Parkinson’s Disease. Acta Neuropathol. 2013, 125, 795–813. 10.1007/s00401-013-1114-9.

(60) APEX3; Bruker AXS LLC: Madison, WI, USA, 2020.

(61) SAINT-NT Software Reference Manual, Version 4.0; Bruker AXS, Inc.: Madison, WI, 1996.

(62) Sheldrick, G. M. SHELXT – Integrated Space-Group and Crystal-Structure Determination. Acta Crystallogr. Sect. A Found. Crystallogr. 2015, 71, 3–8. 10.1107/S2053273314026370.

(63) Sheldrick, G. M. Crystal Structure Refinement with SHELXL. Acta Crystallogr. Sect. C Cryst. Struct. Commun. 2015, 71, 3–8. 10.1107/S2053229614024218.

(64) Schmidbaur, H.; Schier, A. Argentophilic Interactions. Angew. Chem. Int. Ed. 2015, 54, 746–784. 10.1002/anie.201405936.

(65) Emashova, S. K.; Titov, A. A.; Smol’yakov, A. F.; Chernyadyev, A. Y.; Godovikov, I. A.; Godovikova, M. I.; Dorovatovskii, P. V.; Korlykov, A. A.; Filippov, O. A.; Shubina, E. S. Emissive silver(I) cyclic trinuclear complexes with aromatic amine donor pyrazolate derivatives: way to efficiency. Inorg. Chem. Front. 2022, 9, 5624–5634. 10.1039/d2qi01648f.

(66) Riss, T. L.; Moravec, R. A.; Niles, A. L.; Duellman, S.; Benink, H. A.; Worzella, T. J.; Minor, L. Cell Viability Assays in The Assay Guidance Manual, U.S. National Library of Medicine, 2013.

(67) Wang, X.; Becker, K.; Levine, N.; Zhang, M.; Lieberman, A. P.; Moore, D. J.; Ma, J. Pathogenic alpha-synuclein aggregates preferentially bind to mitochondria and affect cellular respiration. Acta Neuropathol. Commun. 2019, 7:41. 10.1186/s40478-019-0696-4.

(68) Son, H. J.; Kim, S.; Kim, S.-Y.; Jung, J. H.; Lee, S. H.; Kim, S.-J.; Kim, C.; Hahn, A. Three-Dimensional β-Amyloid Burden Correlation Between the Eye and Brain in Alzheimer’s Didease Mice Using Light-Sheet Fluorescence Microscopy. Invest. Ophthalmol. Vis. Sci. 2025, 66, 34. 10.1167/iovs.66.3.34.

(69) Shirota, M.; Ishida, T.; Kinoshita, K. Effects of surface-to-volume ratio of proteins on hydrophilic residues: Decreas in occurrence and increase in buried fraction. Protein Sci. 2008, 17, 1596–1602. 10.1110/ps.035592.108.

(70) Deeg, A. A.; Reiner, A. M.; Schmidt, F.; Schueder, F.; Ryazanov, S.; Ruf, V. C.; Giller, K.; Becker, S.; Leonov, A.; Griesinger, C.;, et al. Anle138b and Related Compounds Are Aggregation Specific Fluorescence Markers and Reveal High Affinity Binding to α-Synuclein Aggregates. Biochim. Biophys. Acta BBA - Gen. Subj. 2015, 1850, 1884–1890. 10.1016/j.bbagen.2015.05.021.

(71) Bungeroth, M.; Schlösser, N.; Frieg, B.; Högen, T.; Tatenhorst, L.; Kordasiewicz, H. B.; Bähr, M.; Urlaub, H.; Zweckstetter, M.; Triller, A.;, et al. Differential Aggregation Properties of Alpha-Synuclein Isoforms. Neurobiol. Aging 2014, 35, 1913–1919. 10.1016/j.neurobiolaging.2014.02.009.

(72) Levin, J.; Giese, A.; Boetzel, K.; Israel, L.; Högen, T.; Nübling, G.; Kretzschmar, H.; Lorenzl, S. Increased α-synuclein aggregation following limited cleavage by certain matrix metalloproteinases. Exp. Neurol. 2009, 215, 201–208. 10.1016/j.expneurol.2008.10.010

(73) Espay, A. J.; Lees, A. J. Loss of Monomeric Alpha-Synuclein (Synucleinopenia) and the Origin of Parkinson’s Disease. Parkinsonism Relat. Disord. 2024, 122, 106077. 10.1016/j.parkreldis.2024.106077.

