## Supplemental for "Coordination of Anle138b to Silver Results in Selective Reduction of a C-terminal truncated Alpha-synuclein Protein and Increased Aggregate Size": Supporting Info (Aug25).docx

**Contents**

Page

Synthesis of 3-(1,3-benzodioxol-5-yl)-1-(3-bromophenyl)-3-hydroxyprop-2-en-1-one…..3

Figure S1. ^1^H NMR spectrum (400 MHz, CDCl_3_, 25 °C) of 3-(1,3-benzodioxol-5-yl)-1-

(3-bromophenyl)-3-hydroxyprop-2-en-1one………………………………………………………………..4

Figure S2. ^13^C NMR spectrum (101 MHz, CDCl_3_), 25 °C) of 3-(1,3-benzodioxol-5-yl)-1-(3-bromophenyl)-3-hydroxyprop-2-en-1-one……………………………………………………………….….4

Synthesis of 5-(1,3-benzodioxol-5-yl)-3-(3-bromophenyl)-1*H*-pyrazole…………………….….5

Figure S3. ^1^H NMR spectrum (400 MHz, CDCl_3_, 25 °C) of 5-(1,3-benzodioxol-5-yl)-3-

(3-bromophenyl)-1*H*pyrazole………………………………………………………………………………………5

Figure S4. ^13^C NMR spectrum (101 MHz, CDCl_3_), 25 °C) of 5-(1,3-benzodioxol-5-yl)-3-(3-bromophenyl)-1*H*-pyrazole…………………………………………………………………………………………6

Disorder Modeling for compounds[Cu(μ-L)]_3_ and [Ag(μ-L)]_3_………………………………..6

Figure S5. Thermal ellipsoid plot (50% probability) of **H_2_L(ClO_4_)**∙(CH_3_)_2_CO

showing the disordered ClO_4_^-^ anion (left) and crystal packing (right)………………….7

Figure S6. Thermal ellipsoid plot (50% probability) of **[Cu(μ-L)]_3_** showing

the disorder of one μ-L (left) and crystal packing (right)………………………………....8

Figure S7. Thermal ellipsoid plot (50% probability) of **[Ag(*μ*-L)]_3_** showing

the disordered μ-L groups (left) and crystal packing (right)………………………………8

Figure S8. UV-vis absorption spectra of **HL**, **H_2_L(ClO_4_)**, and **[Cu(μ-L)]_3_**

in XXXXX, 295 K……………………………………………………………………………...9

Figure S9. Emission spectra of **HL**, **H_2_L(ClO_4_)**, and **[Cu(μ-L)]_3_** in MeOH/thf, 295 K....9

Figure S10. Emission spectrum of solid **[Ag(μ-L)]_3_**, 295 K, upon 350 nm excitation...10

Figure S11. Images of non-differentiated SHSY5Y cells, etc……………………………10

Figure S12. **HL**, **H_2_LClO_4_**, **[Cu^I^(µ-L)]_3_**, and **[Ag^I^(µ-L)]_3_**. Plate images of MTT, etc……11

Figure S13. Edman Sequencing results, etc………………………………………………12

Figure S14. PFFs treatment greatly induces the amount of α-synuclein, etc………….13

Figure S15. Molecular weight distribution of α-synuclein species detected, etc………14

**Synthesis of 3-(1,3-benzodioxol-5-yl)-1-(3-bromophenyl)-3-hydroxyprop-2-en-1-one.**

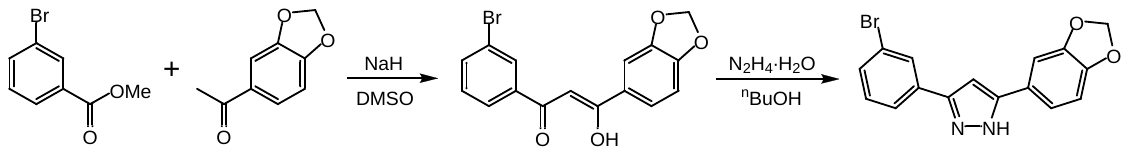

The reaction was carried out under an N_2_ atmosphere in a three-necked flask charged with NaH (~60% dispersion in mineral oil; 2.500 g, 62.50 mmole) and equipped with a pressure-equalizing addition funnel, a reflux condenser and a rubber septum. The mineral oil was washed from the NaH with hexanes (3 ⨯ 50 mL) by using a syringe with a needle inserted through the septum. Dimethyl sulfoxide (40 mL) was added to the flask, which was cooled in an ice-bath. Next, a solution of 1-(1,3-benzodioxol-5-yl)ethanone (8.200 g, 50.00 mmole) and methyl 3-bromobenzoate (13.450 g, 62.50 mmole) in dimethyl sulfoxide (30 mL) was slowly added from the dropping funnel under stirring over 2 hours. The ice-bath was removed, the reaction mixture was stirred at room temperature for 20 hours and then it was carefully poured into an ice-water mixture (250 mL) containing 85% H_3_PO_4_ (2.5 mL) under stirring. After stirring for 1 hour, the precipitate was filtered out, washed with water (2 ⨯ 60 mL) and dried under high vacuum for 12 hours. The crude product was recrystallized from a 1:1 mixture of ethanol and ethyl acetate (200 mL) by cooling the hot solution to room temperature and then keeping it in an ice-bath for 1 hour. The pale yellow crystalline solid was filtered out and dried under high vacuum in a warm water-bath for 12 hours. Yield: 15.471 g (89%). ^1^H NMR (400 MHz, CDCl_3_): 16.79 (s, 1H), 8.06 (t, 1H, ^4^*J* = 2 Hz), 7.86 (d, 1H, ^3^*J* = 8 Hz), 7.64 (d, 1H, ^3^*J* = 8 Hz), 7.59 (dd, 1H, ^3^*J* = 8 Hz, ^4^*J* = 2 Hz), 7.45 (d, 1H, ^4^*J* = 1 Hz), 7.34 (t, 1H, ^3^*J* = 8 Hz), 6.89 (d, 1H, ^3^*J* = 8 Hz), 6.68 (s, 1H), 6.06 (s, 2H) ppm. The enol/diketone tautomeric ratio is approximately 93/7, based on integration of the peaks at 16.79 ppm (enol) and 4.52 ppm (diketone). The chemical shifts shown above correspond to the major tautomer. ^13^C NMR (101 MHz, CDCl_3_): 186.5, 182.1, 151.8, 148.4, 137.5, 135.1, 130.3, 130.1, 129.9, 125.6, 123.3, 123.0, 108.4, 107.4, 102.1, 92.8 ppm.

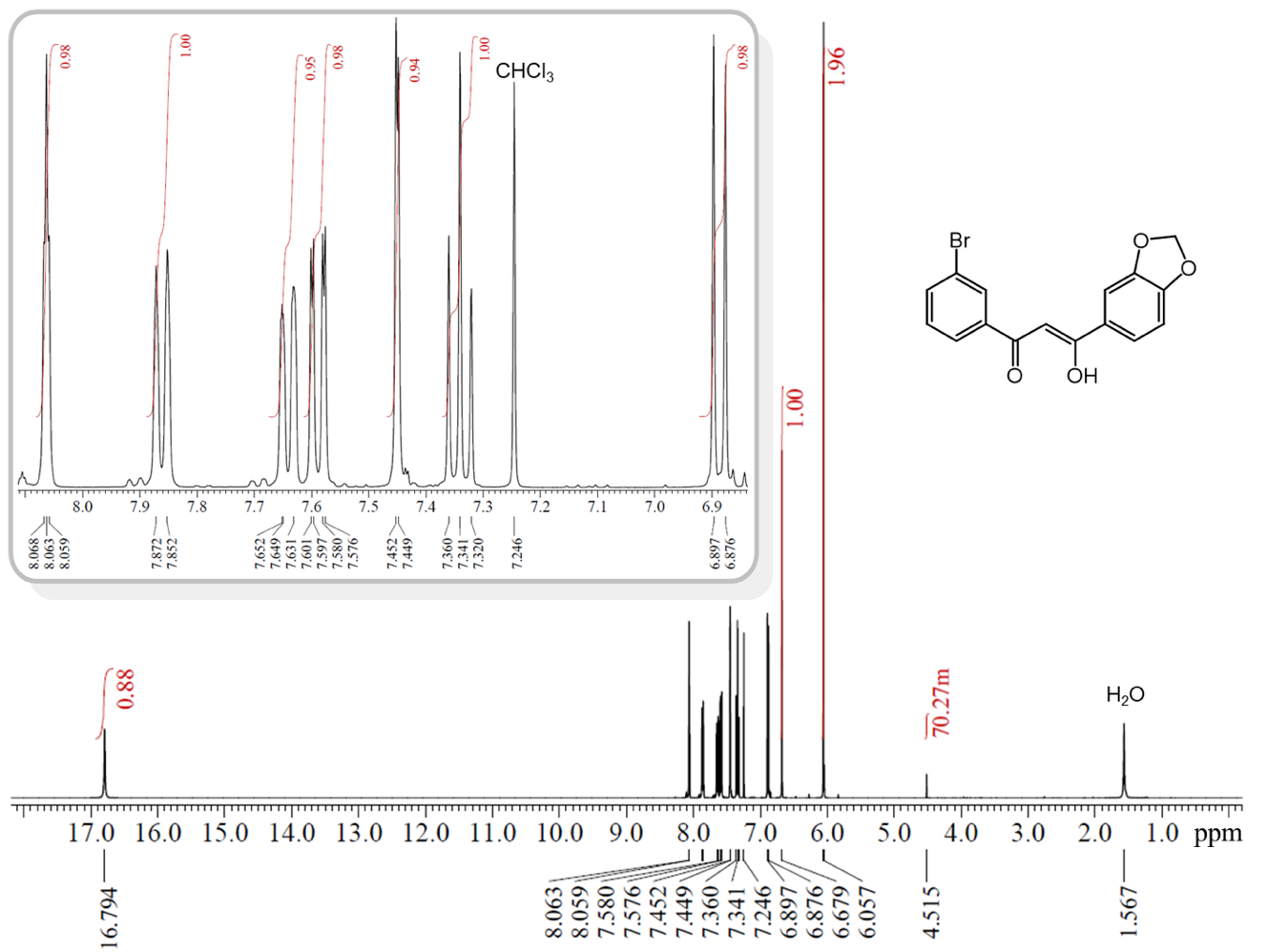

**Figure S1**. ^1^H NMR spectrum (400 MHz, CDCl_3_, 25 °C) of 3-(1,3-benzodioxol-5-yl)-1-(3-bromophenyl)-3-hydroxyprop-2-en-1-one.

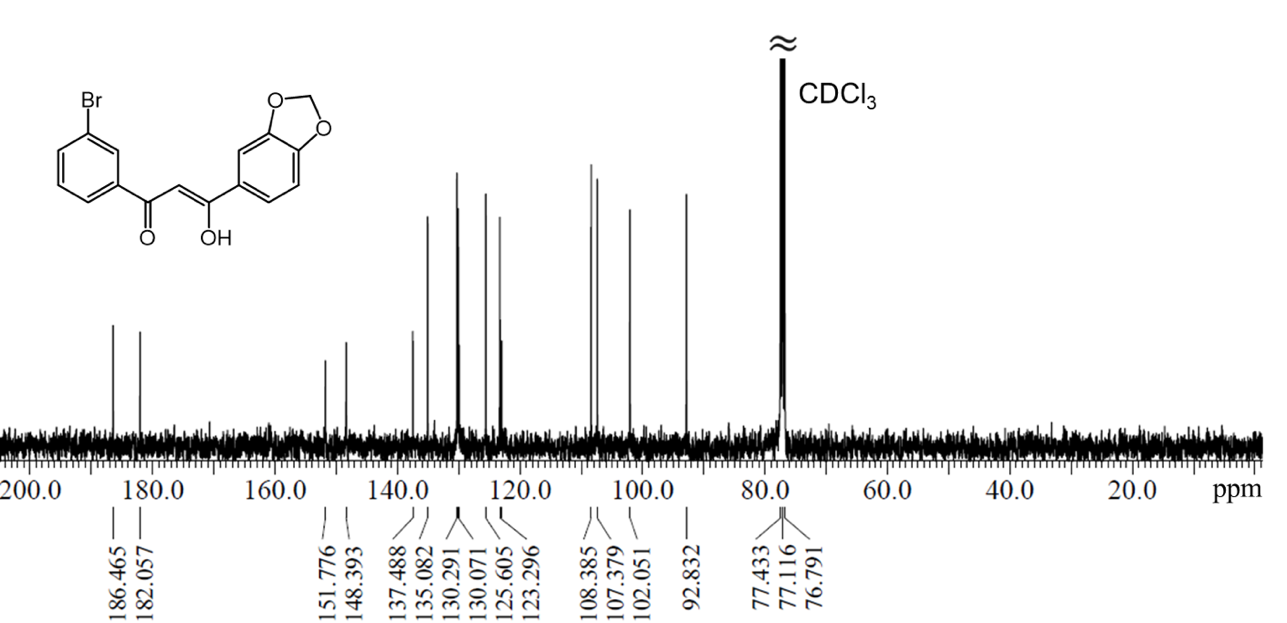

**Figure S2**. ^13^C NMR spectrum (101 MHz, CDCl_3_), 25 °C) of 3-(1,3-benzodioxol-5-yl)-1-(3-bromophenyl)-3-hydroxyprop-2-en-1-one.

**Synthesis of 5-(1,3-benzodioxol-5-yl)-3-(3-bromophenyl)-1*H*-pyrazole**.

3-(1,3-Benzodioxol-5-yl)-1-(3-bromophenyl)-3-hydroxyprop-2-en-1-one (15.471 g, 44.56 mmol) and hydrazine monohydrate (5.20 mL, 5.37 g, 107.2 mmol) were refluxed in *n*-butanol (100 mL) for 4 hours. After cooling to room temperature, the flask was cooled in an ice-bath for 1 hour. The white solid was filtered out, washed with water (250 mL) and dried under high vacuum in a warm water-bath for 12 hours. Yield: 14.300 g (94%). ^1^H NMR (400 MHz, CDCl_3_): 13.30 (s, 1H), 7.98 (s, 1H), 7.79 (d, 1H, ^3^*J* = 8 Hz), 7.48 (d, 1H, ^3^*J* = 8 Hz), 7.37 (t, 1H, ^3^*J* = 8 Hz), 7.35 (s, 1H), 7.29 (dd, 1H, ^3^*J* = 8 Hz, ^4^*J* = 2 Hz), 7.17 (s, 1H), 6.97 (d, 1H, ^3^*J* = 8 Hz), 6.03 (s, 2H) ppm. ^13^C NMR (101 MHz, CDCl_3_): 150.2, 148.3, 147.6, 144.1, 136.5, 131.5, 130.8, 128.0, 124.5, 123.8, 122.8, 119.4, 109.2, 106.1, 101.8, 100.3 ppm. HRMS (ESI-TOF) *m/z*: [M‒H]^−^ calcd for C_16_H_10_BrN_2_O_2_ 340.9926, found 340.9955.

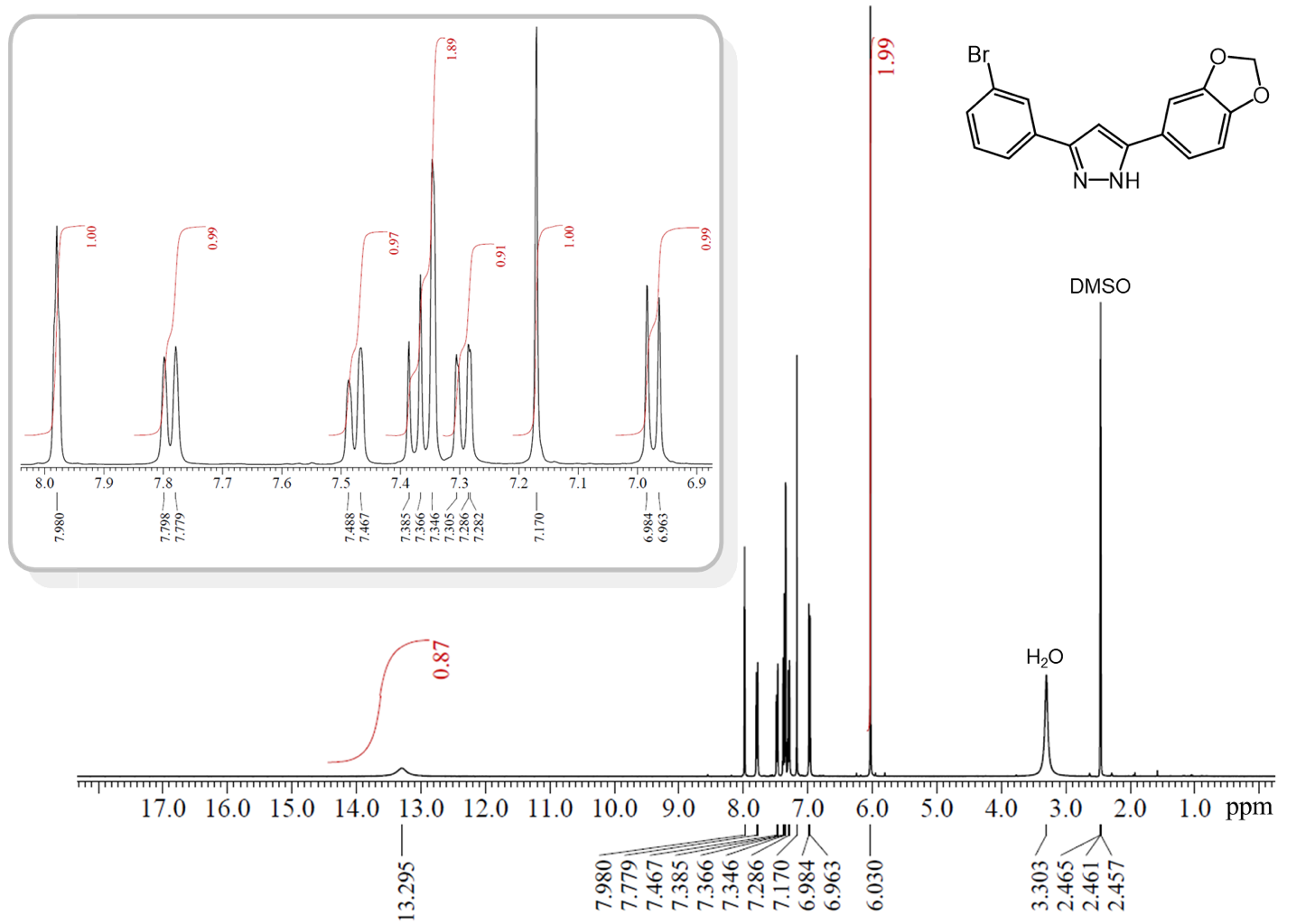

**Figure S3**. ^1^H NMR spectrum (400 MHz, CDCl_3_, 25 °C) of 5-(1,3-benzodioxol-5-yl)-3-(3-bromophenyl)-1*H*-pyrazole.

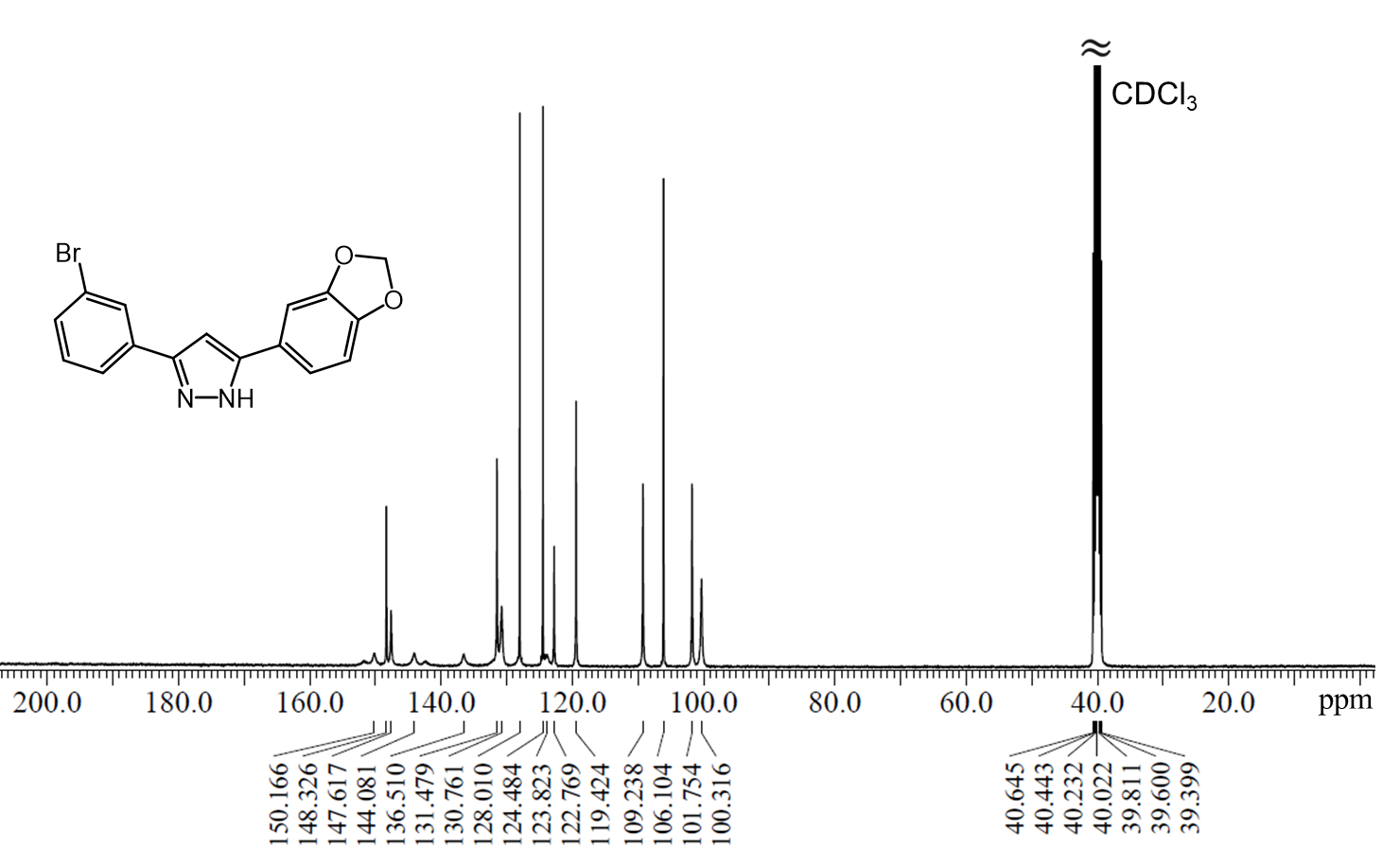

**Figure S4**. ^13^C NMR spectrum (101 MHz, CDCl_3_), 25 °C) of 5-(1,3-benzodioxol-5-yl)-3-(3-bromophenyl)-1*H*-pyrazole.

**Disorder Modeling for compounds[Cu(μ-L)]_3_ and [Ag(μ-L)]_3_**

Upon initial refinement, electron density analysis indicated disorder involving two bromophenyl rings. To account for this, a two-component disorder model was implemented. The occupancy factors for the two disordered components of the bromophenyl rings were assigned as **21** and **-21**, representing occupancies of 0.50 and 0.50, respectively, within a free variable system. The atoms corresponding to these disordered components were explicitly defined using **PART 1** and **PART 2** commands in the input file. An intermediate refinement cycle was performed to allow the program to optimize the positions and thermal parameters of these initial disordered components.

Following the intermediate refinement, the difference Fourier map revealed clear electron density corresponding to additional missing five-membered rings. These atoms were subsequently added to the model and assigned to the **other PART** of the disorder model, consistent with their observed electron density. Furthermore, the phenyl rings attached to the pyrazole were also included in this disorder model. Importantly, the pyrazole ring itself remained fixed as **PART 0**, indicating no observed disorder.

To achieve a stable and chemically reasonable refinement of the disordered fragments, specific restraints and constraints were applied. The geometric parameters of the disordered fragments were restrained using either **SAME/ADI** commands. **SAME** was applied where chemically identical fragments were constrained to possess identical bond lengths and angles, requiring careful attention to atom ordering. Alternatively, **SADI** was used to restrain equivalent bond lengths to be similar, where slight conformational differences were plausible. To ensure physical realism of the atomic displacement parameters (ADPs) and prevent non-positive definite ellipsoids, **SIMU 0.01** commands were applied to all disordered atoms. This constraint encourages neighboring disordered atoms to adopt similar thermal parameters.

With the comprehensive disorder model and associated restraints in place, the refinement proceeded through iterative cycles. The refinement was monitored for convergence, chemical plausibility of bond lengths and angles, and the absence of significant residual electron density or negative ADPs, ultimately leading to a reliable refinement.

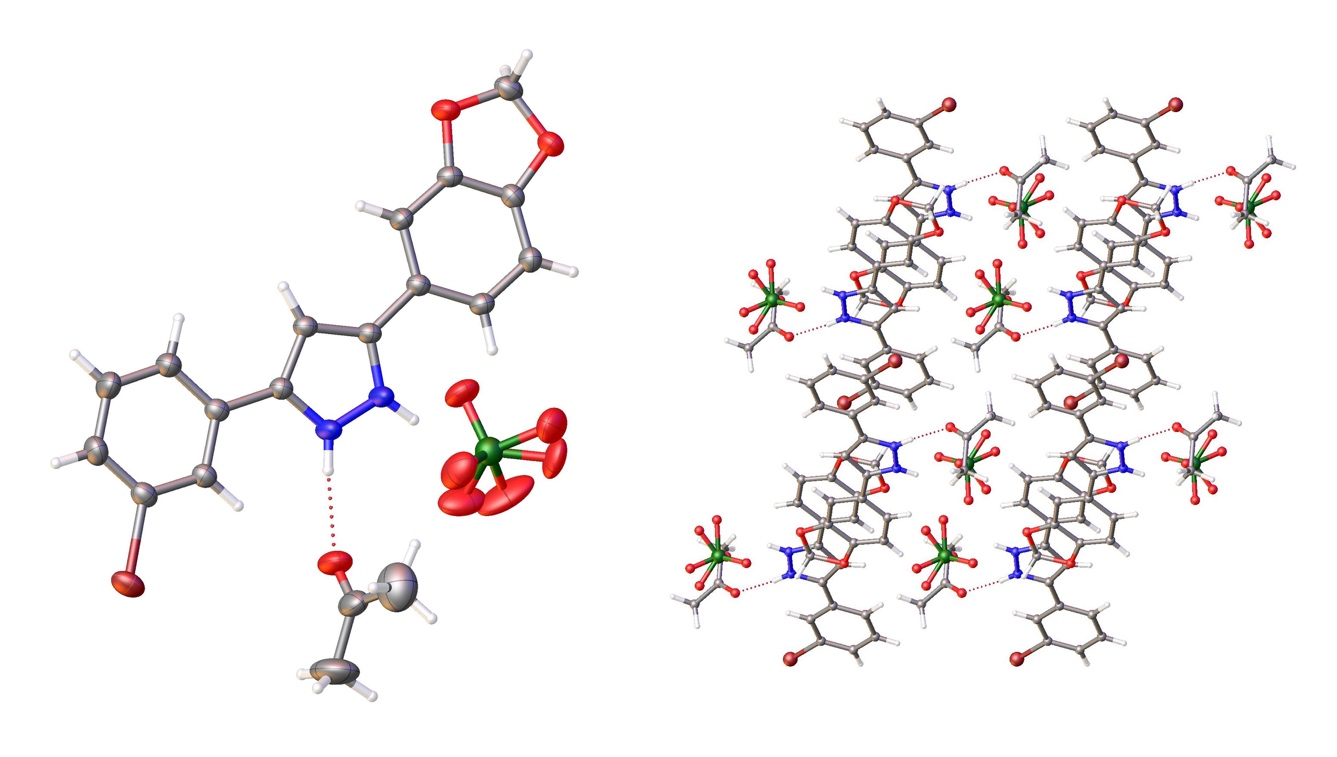

**Figure S5**. Thermal ellipsoid plot (50% probability) of **H_2_L(ClO_4_)**∙(CH_3_)_2_CO showing the disordered ClO_4_^-^ anion (left) and crystal packing (right).

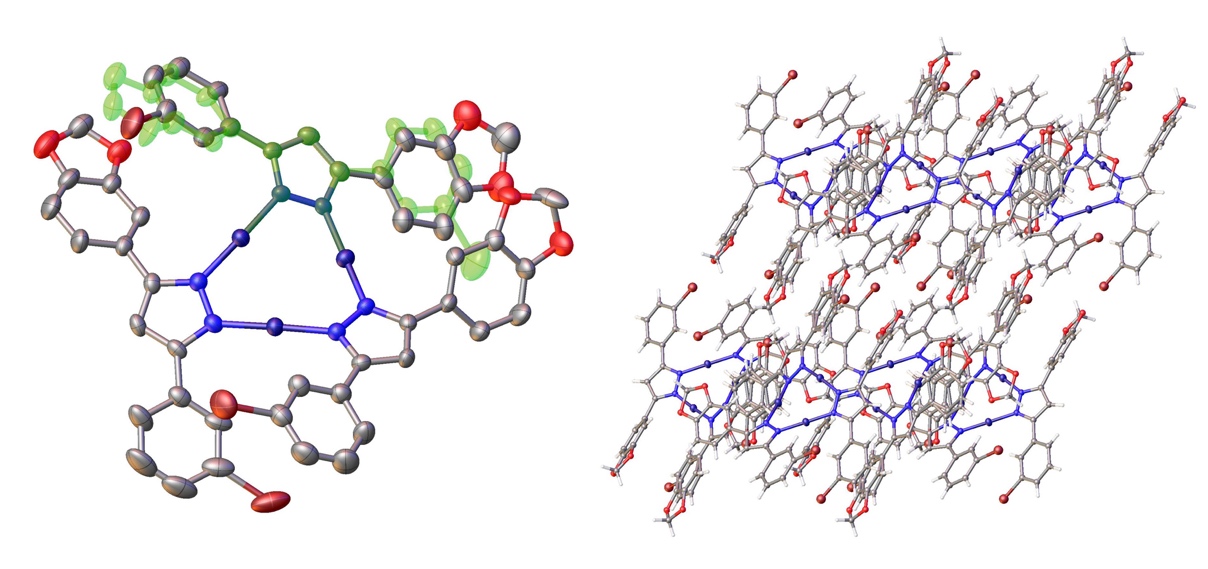

Figure S6. Thermal ellipsoid plot (50% probability) of **[Cu(μ-L)]_3_** showing the disorder of one μ-L (left) and crystal packing (right).

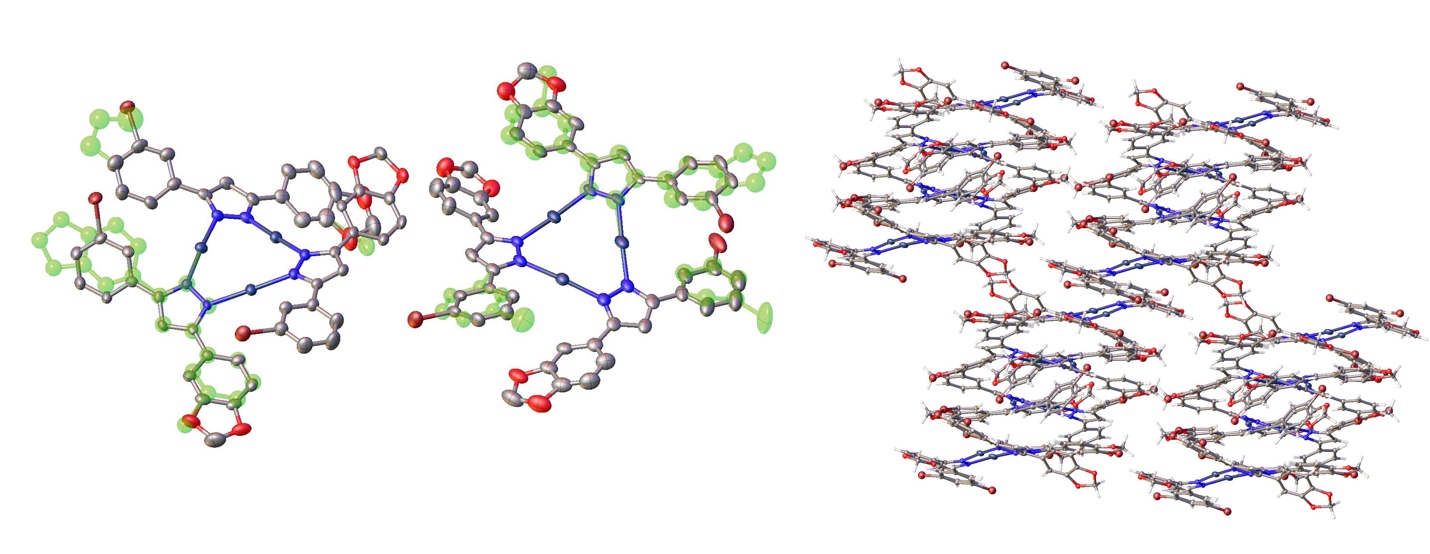

Figure S7. Thermal ellipsoid plot (50% probability) of **[Ag(*μ*-L)]_3_** showing the disordered μ-L groups (left) and crystal packing (right).

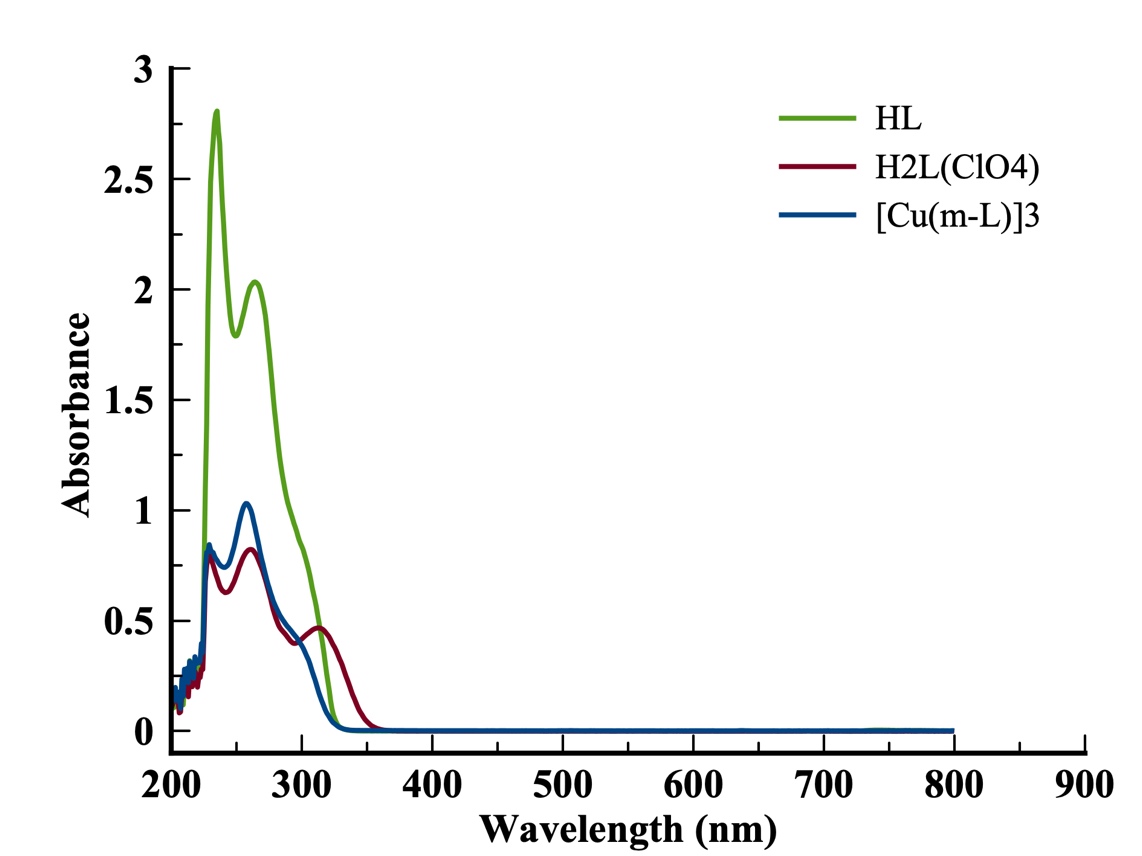

Figure S8. UV-vis absorption spectra of **HL**, **H_2_L(ClO_4_)**, and **[Cu(μ-L)]_3_** in XXXXX, 295 K.

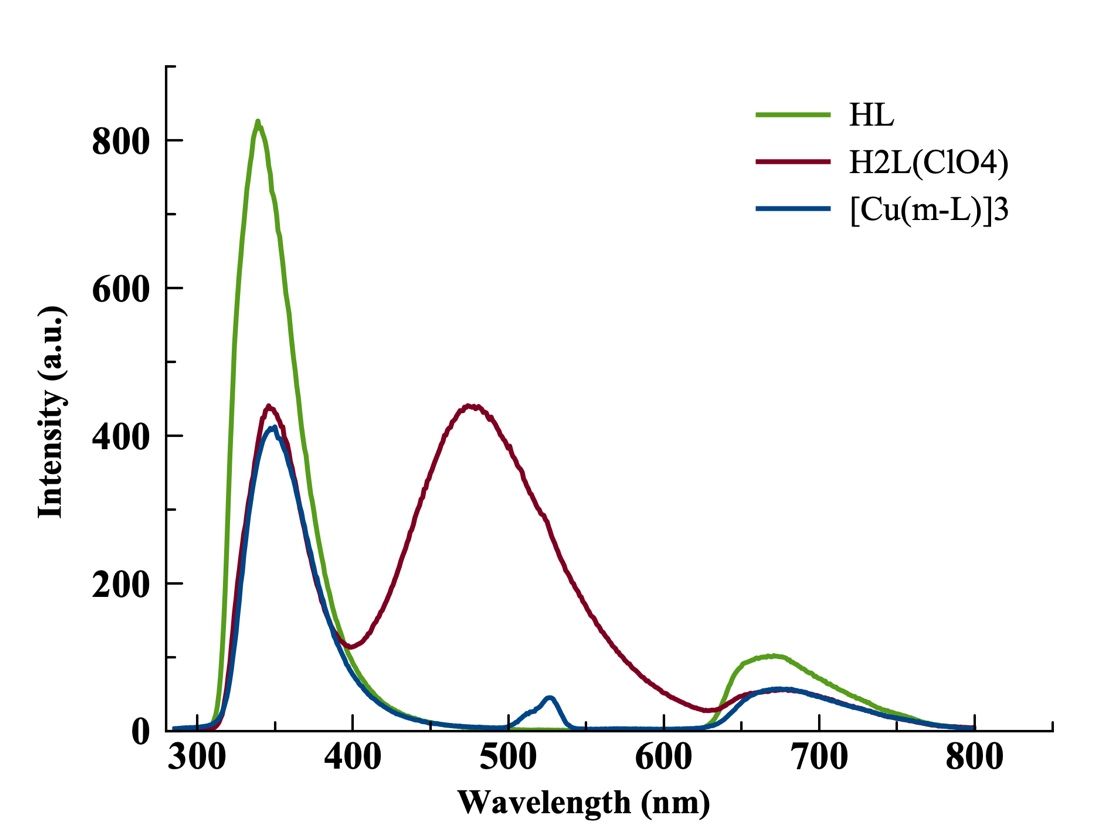

Figure S9. Emission spectra of **HL**, **H_2_L(ClO_4_)**, and **[Cu(μ-L)]_3_** in MeOH/thf, 295 K.

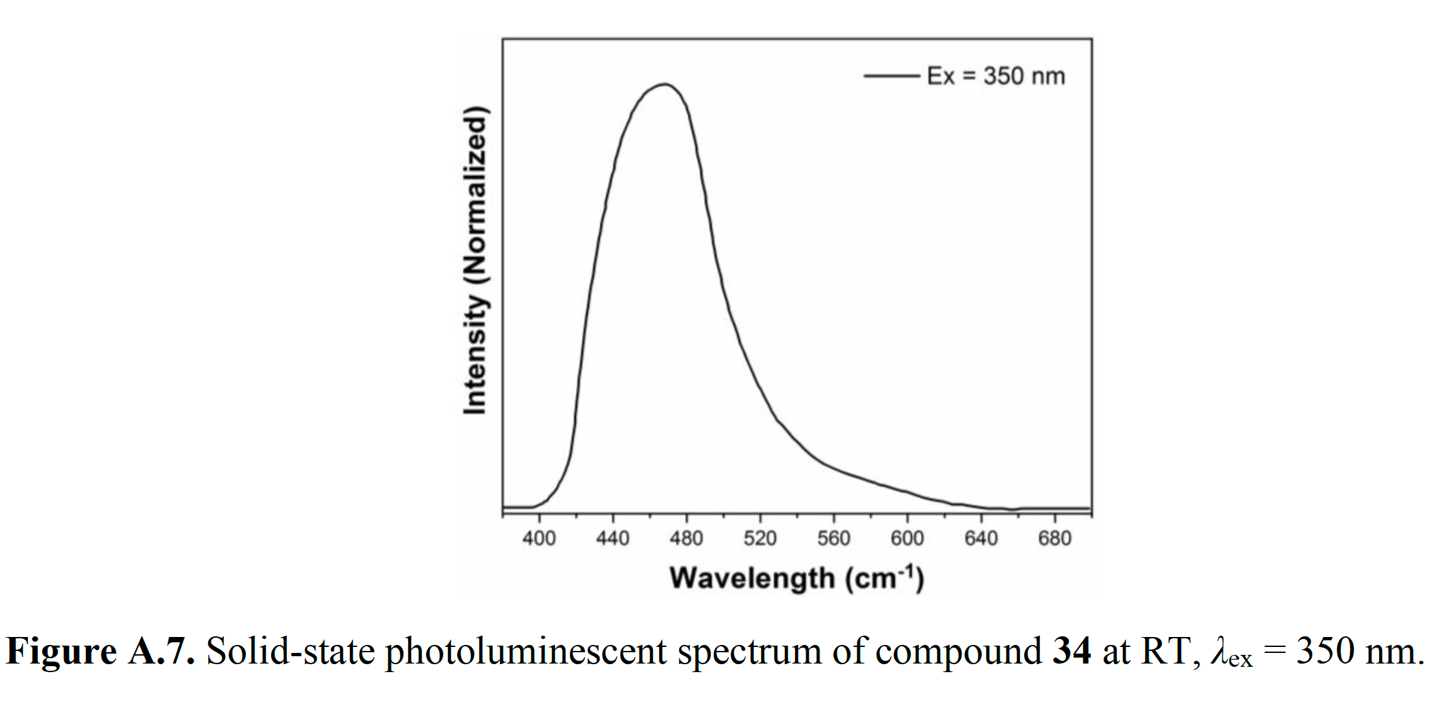

Figure S10. Emission spectrum of solid **[Ag(μ-L)]_3_**, 295 K, upon 350 nm excitation.

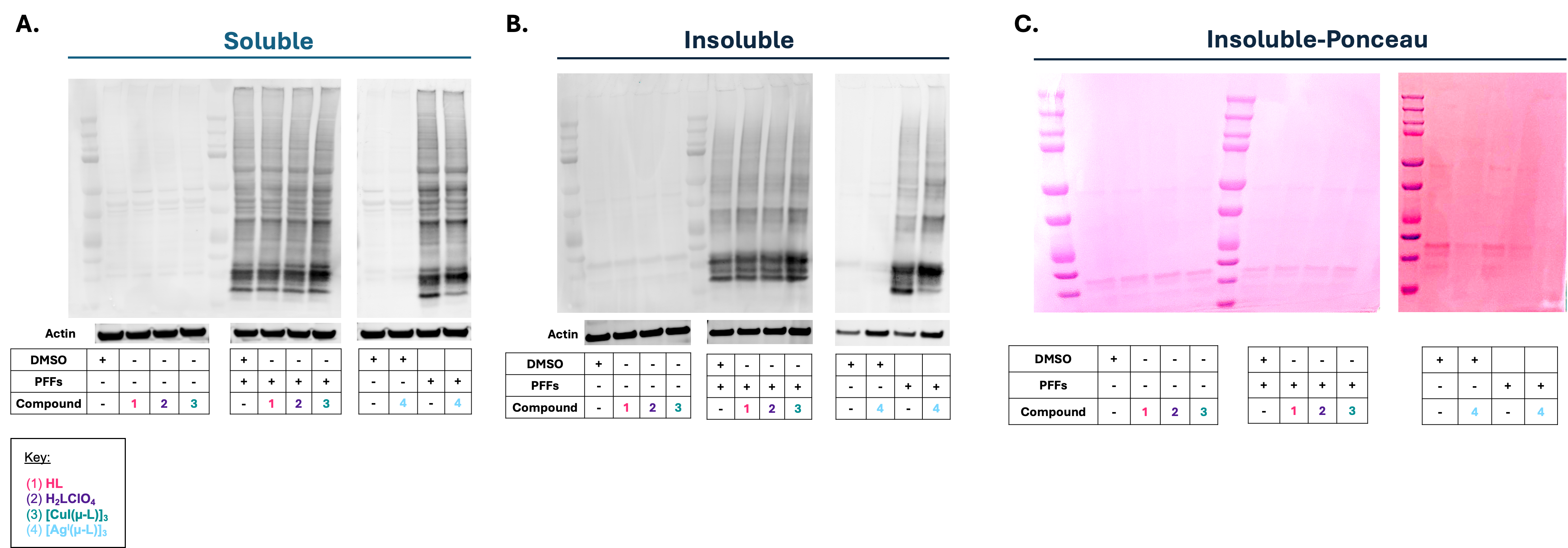

Figure S11. PFFs treatment greatly induces the amount of α-synuclein signal detected with BD antibody in 48 hours in the soluble **(A)** and insoluble **(B)** fractions as compared to control. **(C)** Ponceau S staining for the insoluble fraction shown on the right, indicating similar protein loading in soluble and insoluble fractions.

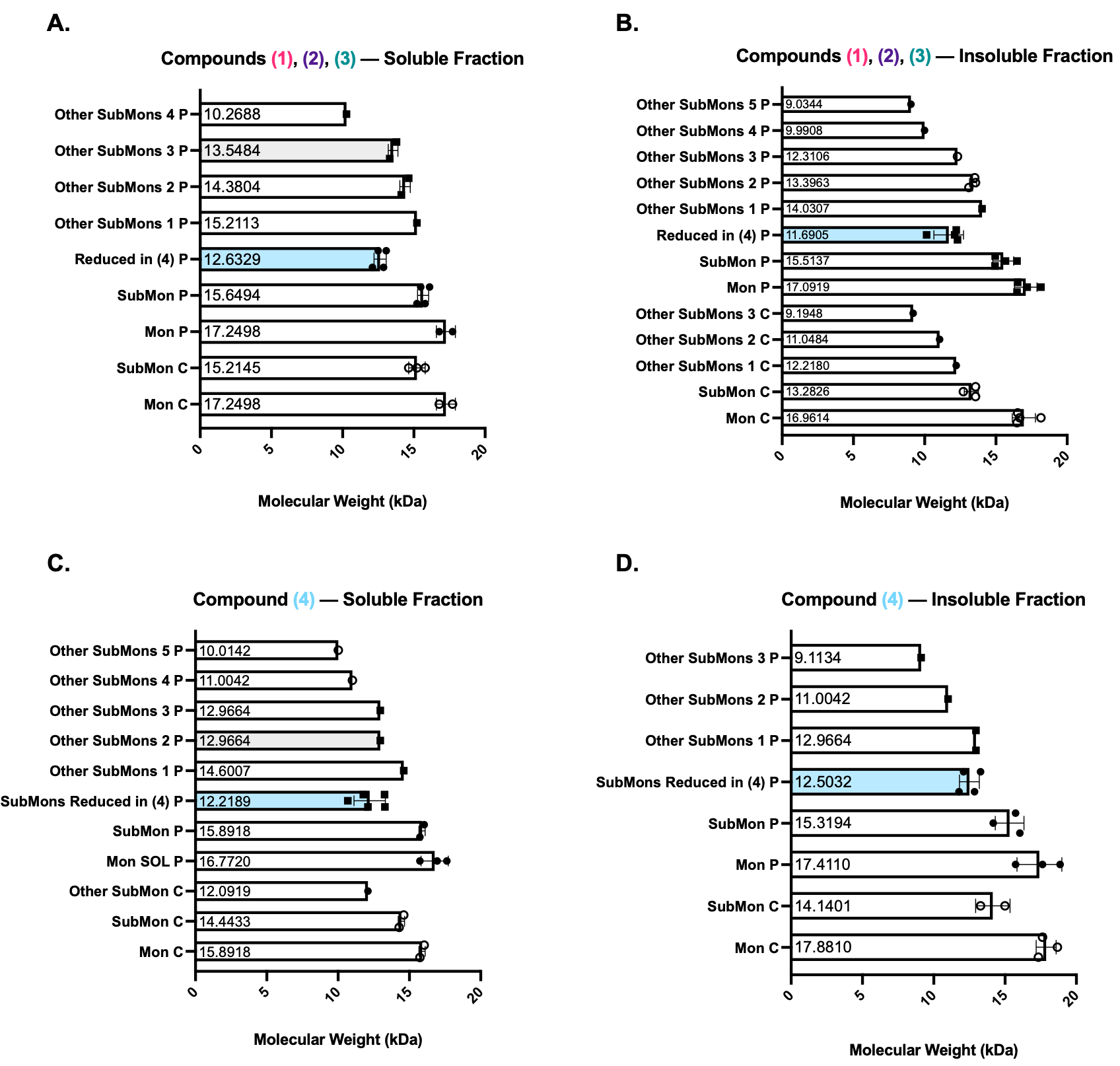

Figure S12. Molecular weight distribution of α-synuclein species detected by BD antibody were calculated using ImageJ from all biological replicates where bands were visible (each dot represents a replicate) for each compound: **HL**, **H_2_L(ClO_4_)**, **[Cu(μ-L)]_3_** and **[Ag(μ-L)]_3_** (labeled 1, 2, 3, and 4, respectively) in the soluble **(A, C)** and in the insoluble **(B, D)** fractions. The band reduced by **[Ag(μ-L)]_3_** treatment is the ~12.4 kDa band is highlighted in blue **(C,D)**.

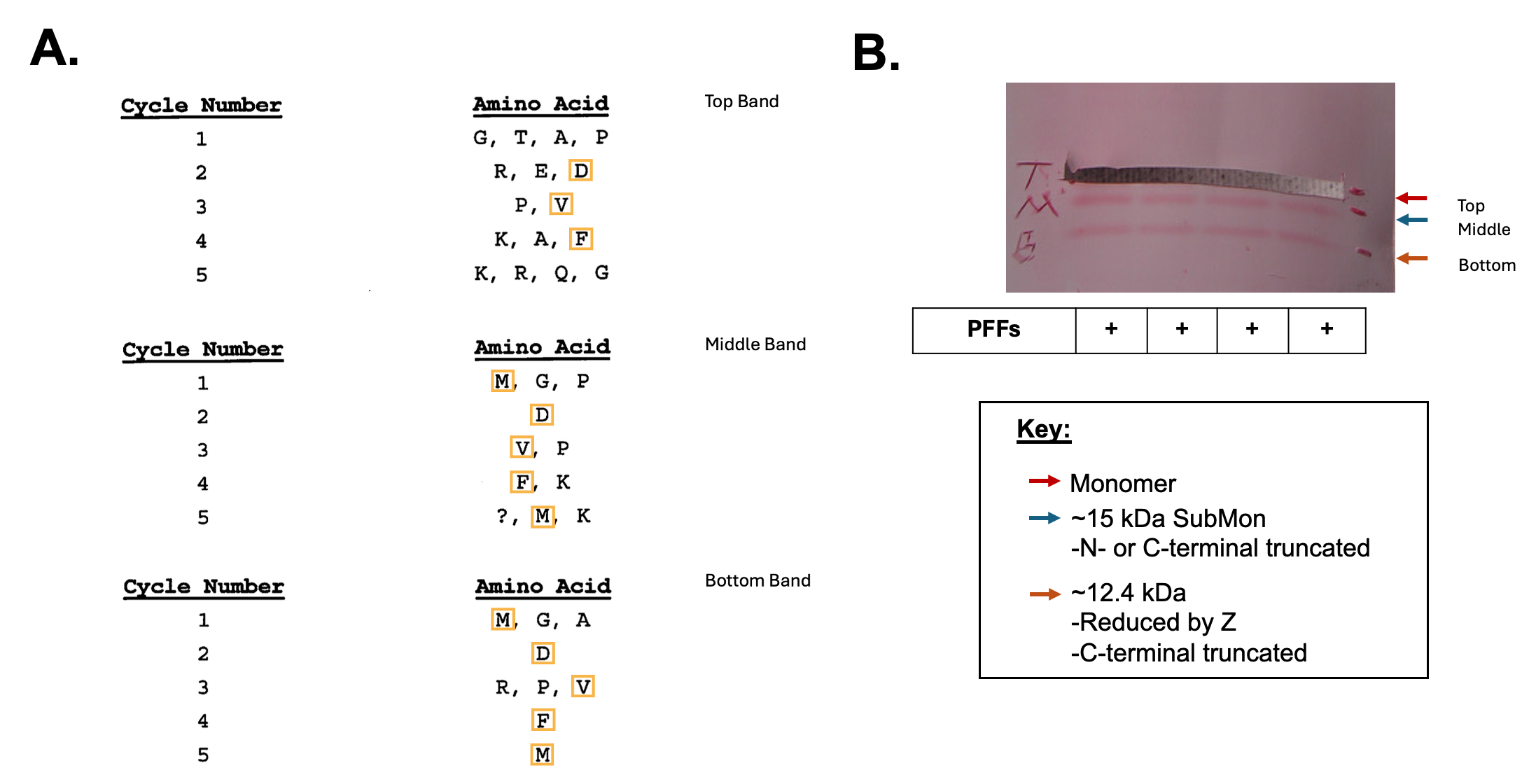

Figure S13. Edman Sequencing results of three bands (indicated by ‘top’, ‘middle’, and ‘bottom’ labels) in PFFs treated non-differentiated SHSY5Y cells from the insoluble fraction. **(A)** Results suggest that the submonomeric species (middle and bottom bands) are C- and not N-terminally truncated as determined by the presence of the MDVFM motif — the first five amino acids of the N-terminus — in the alpha-synuclein protein sequence (yellow squares.) The first amino acid listed in each cycle showed the strongest signal. **(B)** Ponceau S-stained blot loaded onto the sequencer with the top band excised. Four technical replicates were pooled together for each band i.e. top, middle, or bottom in order to increase signal strength. While the exact identity of the bands cannot be determined with this staining method, based on patterns observed earlier, the ‘top’ band is likely the monomer (red arrow), while the middle band is likely the 15 kDa (blue arrow.) The bottom band is hypothesized to be the ~12.4 or 13 kDa C-terminally truncated alpha-synuclein protein (n=4 technical replicates).

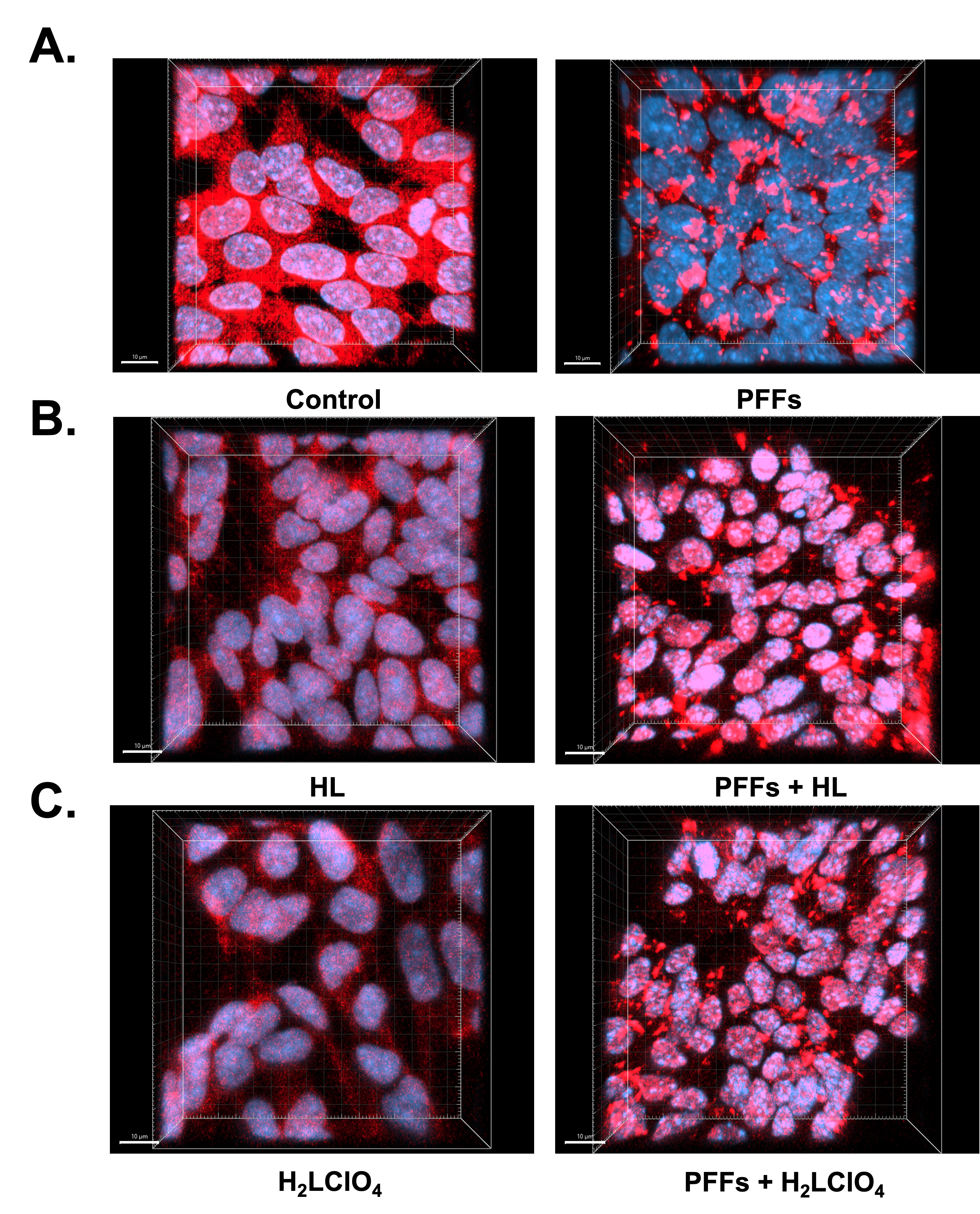

Figure S14. Representative confocal images of SHSY5Y cells show increased alpha-synuclein aggregation (red) with PFFs addition and decrease in puncta with **HL** and **H_2_LClO_4_** addition**.**

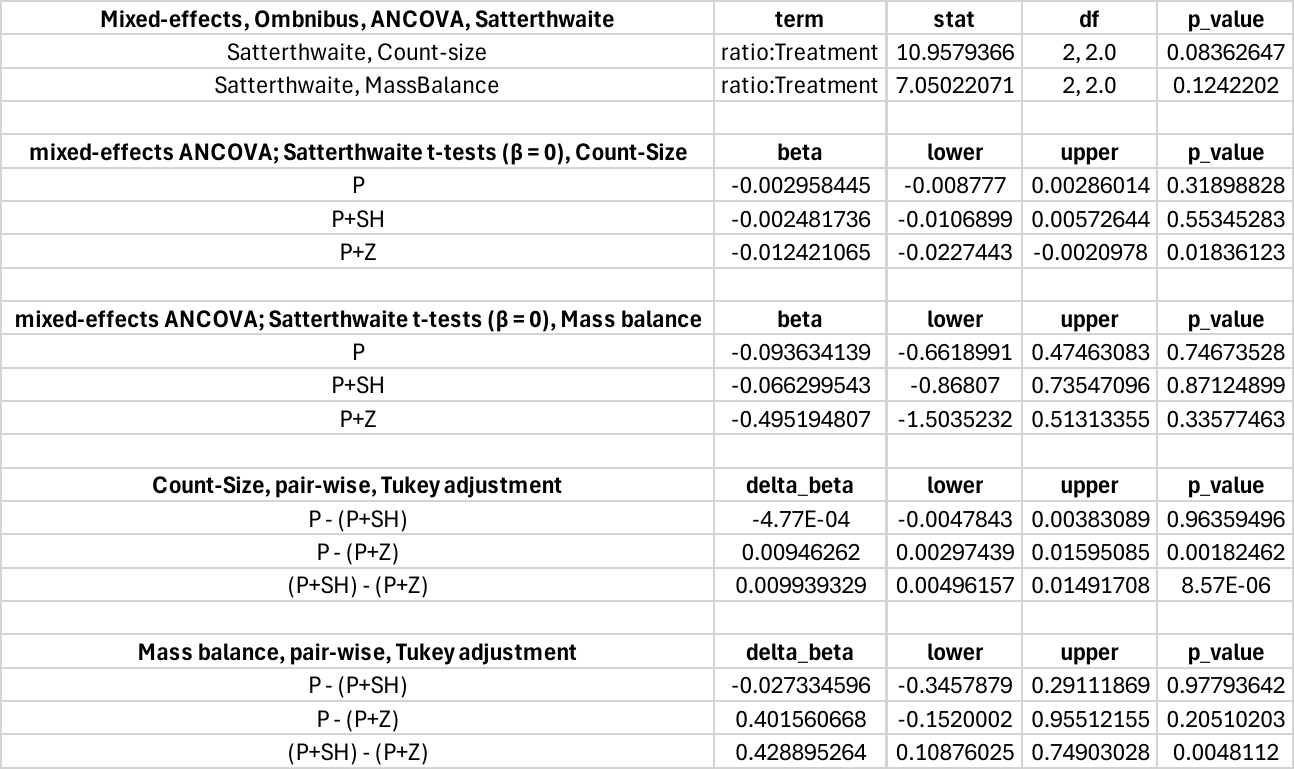

Table 1. **[Ag^I^(**µ**-L)]_3_** has larger puncta as compared to a smaller number of puncta/nuclei. The omnibus interaction shows a trend: β = -0.012, 95% CI [-0.023, -0.002], Satterthwaite comparison, (F(2,2.0)=10.96, p=0.084). Pair-wise slope contrasts confirm **[Ag^I^(**µ**-L)]_3_** < P (Δβ = −0.0095 [−0.016, −0.003], p<0.0018) and **[Ag^I^(**µ**-L)]_3_** < **[Cu^I^(**µ**-L)]_3_** (Δβ = −0.001 [−0.015, −0.005], p<0.0001); P and **[Cu^I^(**µ**-L)]_3_** do not differ (p=0.96.] **[Ag^I^(**µ**-L)]_3_** omnibus interaction shows a trend towards larger puncta mass per a smaller number of puncta/nuclei for **[Ag^I^(**µ**-L)]_3_** (β = -0.50, 95% CI [-1.50, 0.51], Satterthwaite, F(2,2.0)=7.05, p=0.12.) Pair-wise slope contrasts show **[Ag^I^(**µ**-L)]_3_** is steeper than **[Cu^I^(**µ**-L)]_3_** (Δβ = −0.43 [−0.75, −0.11], p<0.0048), while difference vs P is not significant for **[Ag^I^(**µ**-L)]_3_** (p=0.21) nor **[Cu^I^(**µ**-L)]_3_** (p=0.98.)
